# Dystrophin Dp71 is essential for the development and function of macrophages

**DOI:** 10.1101/2025.04.24.650386

**Authors:** Natalia Chira, Julian Swatler, Antigoni Manousopoulou, Robin Rumney, Jacek Hajto, Aleksandra Oksiejuk, Diana Garay Baquero, Justyna Rog, Cory H. White, Stefan Hinz, Nancy Alnassar, Christopher Young, Stephen Arkle, Michal Korostynski, Ewa Kozłowska, Spiros D. Garbis, Dariusz C. Górecki

## Abstract

Mutations in the *DMD* gene, encoding dystrophins, cause progressive muscle degeneration with severe sterile inflammation. While macrophages predominate amongst muscle-infiltrating cells, being central to both damage and regeneration, they were not known to express dystrophin. Yet, we recently demonstrated Dp71 dystrophin expression correlating with tumour infiltrating macrophages. Here we report physiological, developmentally regulated expression of Dp71 in human and mouse hematopoietic stem cells, which decreases with cell maturation into bone marrow macrophages (BMM). Proteomics with molecular and functional analyses in mouse dystrophin-null BMM and peritoneal macrophages reveal that absence of dystrophin disturbs their development. Alterations in over 300 proteins mapped to pathways and networks relating to reduced migration and phagocytosis and increased NLRP3 inflammasome functions. These defects are Dp71-dependent and not caused by the dystrophic environment, since *Dmd*^mdx^ mouse macrophages, which express Dp71, are not affected. Thus, we identify an important new role for the *DMD* gene. Altered Dp71 expression in tumour microenvironment cells and in dystrophin-null patients should be investigated to understand the commonalities between DMD and tumours, and potentially identify new treatments.

## Introduction

Mutations in the DMD gene^1,2^ cause Duchenne muscular dystrophy (DMD), a debilitating and incurable neuromuscular disorder. Three full-length 14 kb transcripts encode 427 kDa proteins, and further intragenic promoters drive expression of truncated variants, with specific spatiotemporal distribution patterns and tissue functions. While the muscle dystrophy is the main pathology resulting from the loss of dystrophin expression, there is growing evidence that *DMD* gene mutations produce a range of significant cell-autonomous abnormalities in non-muscle cells^3^. The array of impaired functions ranges from brain cells, where it manifests in neuro-behavioural deficits^4,5^, through platelet aggregation defects^6^ to various tumours^7^, where recent studies found differential *DMD* gene expression associated with Duchenne-like abnormalities^8–10^. Moreover, there is ample evidence from both human and animal models that DMD pathology starts already during the development^11–17^.

Interestingly, until recently, little attention was given to the role of dystrophin in the immune cells, despite the well-documented evidence of their importance in the initiation and progression of the dystrophic phenotype^18,19^. Functional alterations in both leukocyte recruitment^20^ and in effector cells from DMD patients^21,22^ have been described. Importantly, inflammation has been shown to precede the dystrophic muscle damage^23–25^. DMD-associated breakdown in the peripheral tolerance has also been described, where adoptive transfer of primed immune cells resulted in transmission of pathology to healthy mice^26^. Treatments reducing immune cell infiltrations also significantly improved the dystrophic phenotype in patients and in the *Dmd*^mdx^ mouse model^23,24,26–34^.

All these findings made us question whether the dystrophic inflammation is just the response to muscle damage and death or if it could be exacerbated by primary immune cell defects resulting from the alterations of DMD transcripts expression.

The dystrophic mouse muscle contains 20 times more macrophages than the healthy one^35^. Given the prominent role that macrophages have in the initiation and maintenance of inflammation, while also being essential for muscle regeneration^36^, we investigated whether DMD macrophages show autonomous alterations due to *Dmd* gene mutations, which affect their functional behaviour.

Using in-depth discovery proteomics with molecular and functional approaches, we explored developing and mature macrophages from two DMD mouse strains. The *Dmd*^mdx^ (mdx) mice, the most widely used model, have a mutation causing the loss of expression of full-length dystrophins only. This is the most common mutation in DMD patients and therefore little attention has been given to the pathology in rare dystrophin-null cases. However, such mutations are invariably associated with severe cases with neuropsychological symptoms^37^ and an exacerbated muscle phenotype have been described in both patients^38^ and dystrophin-null *Dmd*^mdxβgeo^ (mdx^βgeo^) mice, where altered macrophage infiltration patterns were also identified^39^. Furthermore, we recently demonstrated Dp71 dystrophin expression correlating with tumour infiltrating macrophages^9^. Therefore, cells from this model^40^ were studied.

We found that both human and mouse hematopoietic stem cells (HSC) express transcripts encoding Dp71 dystrophin and that the loss of DMD gene expression during macrophage development from HSC triggers major and persistent functional abnormalities in mature cells.

## Results

### Expression and developmental regulation of DMD gene in macrophages

Given the complexity of the gene, with its multiple promoters (Figure 1) and a complex splicing pattern of mRNAs, we amplified the key regions of *Dmd* transcripts expressed in wild type (WT) mouse peritoneal macrophages (PMϕ). Primers complementary to exons 8 and 9, which are present in the full-length transcripts encoding dystrophins Dp427 and to regions upstream of exon 62 (present in Dp427, Dp260, Dp140 and Dp116 transcripts) yield no amplification. In contrast, Dp71 expression was clearly detectable (Figure 1). No amplification in PMϕ from mdx^βgeo^ dystrophin-null macrophages confirmed specificity of this expression. Dp71 undergoes extensive alternative splicing^41–43^ and these variants expressed in specific cells have been found important for different processes, including cell adhesion, division, water homeostasis and nuclear architecture^44^. RT-PCR analysis in PMϕ showed no significant alternative splicing of exons 71, 71-74 or 77 but found that exon 78 could be alternatively spliced in a proportion of transcripts. Therefore, wild-type macrophages can express the 3’-end variant (Figure 1), encoding an alternative C-terminus, which has specific functional properties^10^.

**Figure 1.**
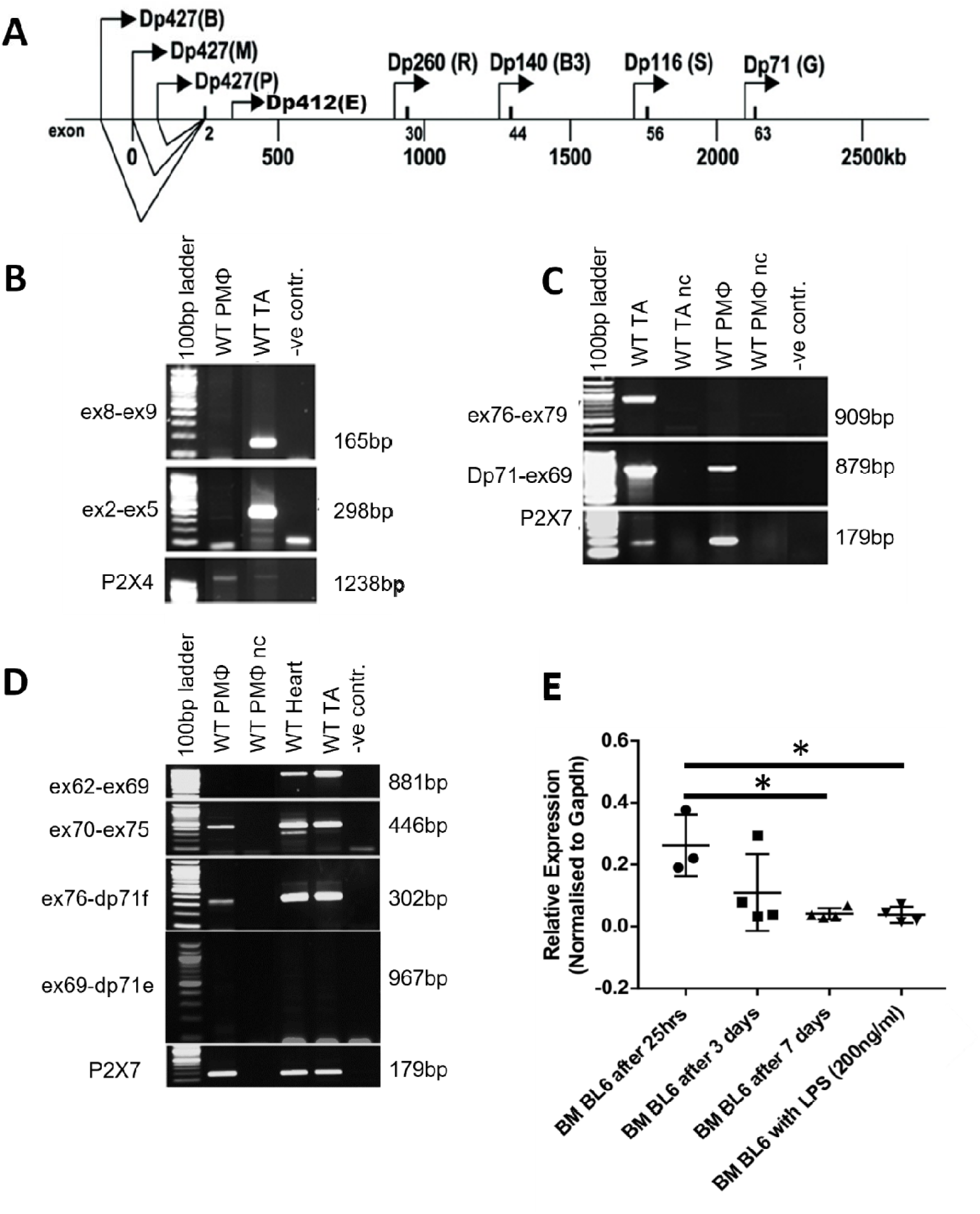
Dmd transcript expression in murine peritoneal macrophages. **(A)** Schematic representation of the human DMD gene, promoters and transcripts. **(B)** Amplification across exons 2 and 5 and exons 8 and 9 (ex8fwd-ex9rev and ex2fwd-ex5rev primers) was used for the identification of the 5’-end of the Dmd transcript. **(C)** Amplification with Dp71fwd-ex69rev primers identified Dp71 isoforms and 76fwd-ex79rev primer pair was used to assess the alternative splicing at the 3’-end. **(D)** Amplification with primers ex62fwd-ex69rev identify transcript other than Dp71; Ex70fwd-ex75rev primer pair identified no alternative splicing of exons 71 or 71-74 in PMϕ while amplification with Ex76fwd-dp71rev pair detected alternative spicing of exon 78. P2X4 and P2X7 primer sets were used as positive cDNA controls, while C57BL/10 tibialis anterior (TA) and heart muscle samples were used as positive controls for Dmd expression. nc denotes samples where the reverse transcriptase was omitted in cDNA synthesis to control for genomic DNA contamination and -ve denotes a no cDNA control. Figure shows representative gel images of at least four replicates. (**E**) qPCR analysis (exon 65-66 primers) of dystrophin expression levels at different stages of BMM development, from bone marrow cells after 25 hrs in culture until day 7, mature macrophages; n ˃ 3, *p < 0.05.

Dp71 is the first dystrophin detectable during organ development, and relatively high Dp71 promoter activity was found associated with morphogenic events and differentiation of several mouse cells^41,45–47^. Therefore, we analysed Dp71 transcript expression at various stages of differentiation of bone marrow-derived macrophages (BMM) *in vitro.* Transcript was found expressed at an early stage and it levels decreased significantly as cells developed into BMM (Figure 1e).

Given the transcript levels significantly decreasing from the early stage of macrophage development, we hypothesised that *Dmd* may play a role even earlier, in hematopoietic stem cells (HSC), and this altered development potentially affects cell functions at subsequent stages, analogous to what was found in muscle development^15^, and in satellite cells^48^.

#### Human and mouse haematopoietic stem cells express dystrophin Dp71

Analysis of the www.bloodspot.eu, the Immunological Genome Project [https://www.immgen.org/] and Cabezas-Wallscheid^49^ expression datasets revealed that both human and mouse HSC express the *DMD* gene (Figure 2a). In mouse HSC and multipotential progenitors (MPP1-4) datasets we found that the signal was obtained from the exons at the 3’-end of the gene only. These detected exons corresponded to the Dp71 transcript. Practically no reads from exons encoding the Dp427 isoforms were identified in any of these transcriptomes (Figure 2b). At the HSC/MPP1 undifferentiated stages the expression levels were at 2.3 and 2.6%, relative to the *Gapdh* housekeeping gene. There was a significant downregulation of dystrophin expression as cells differentiated towards MPP3 (0.9 %) and MPP4 (1 %), i.e., in cells starting to show a myeloid and lymphoid bias, respectively (Figure 2b, Suppl. Figure 1). Thus, Dp71 in hematopoietic cells may contribute to developmental processes and its absence have a downstream functional impact in macrophages that differentiate from such dystrophic HSC.

**Figure 2.**
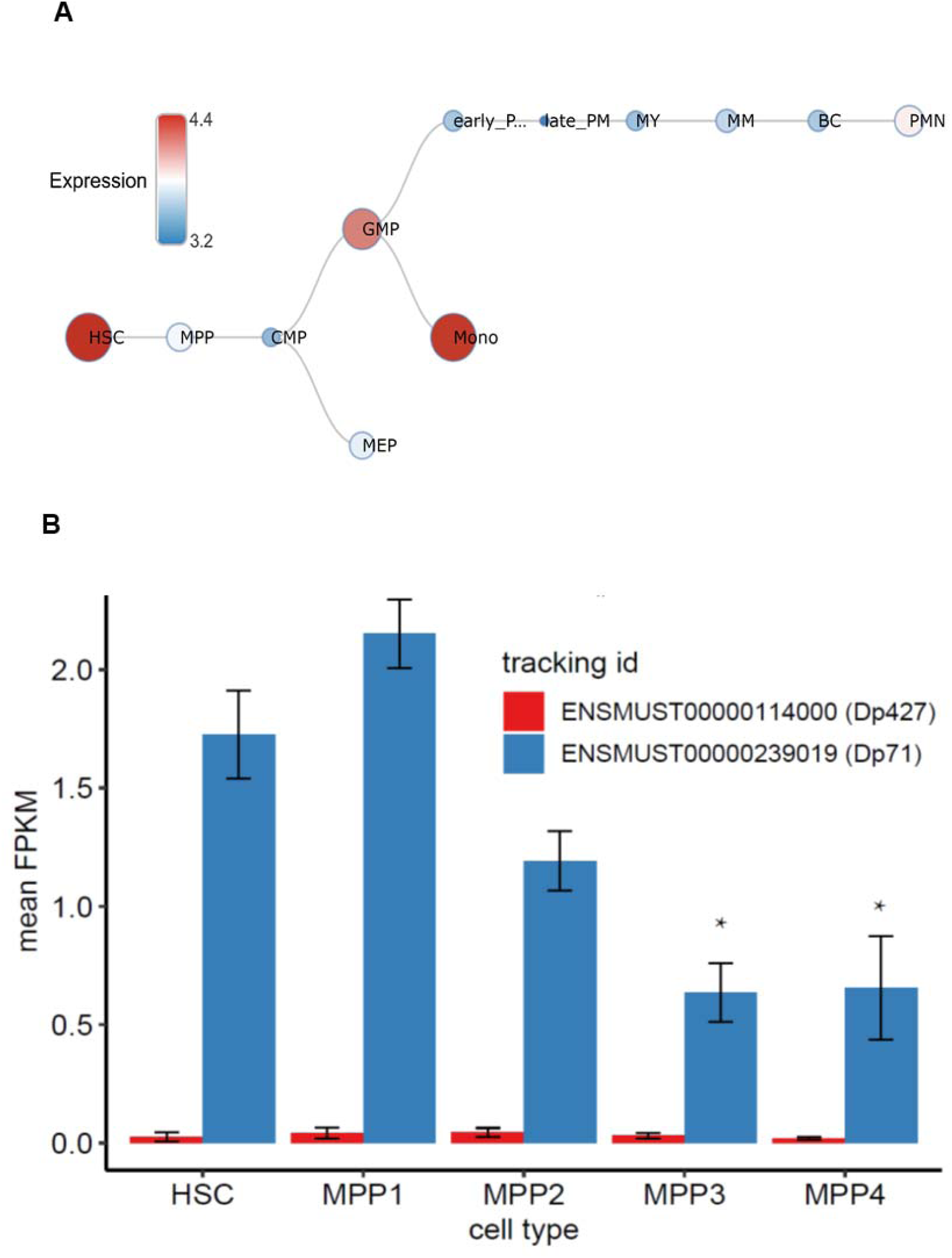
DMD gene expression at different stages of human and mouse HSC development. **A.** Hierarchical differentiation tree. DMD gene expression in human hematopoietic cells at different maturation stages based on curated microarray data from www.bloodspot.eu. HSC: Hematopoietic stem cell (Lin- CD34+ CD38- CD90+ CD45RA-); MPP: Multipotential progenitors (Lin- CD34+ CD38- CD90- 45RA-); CMP: Common myeloid progenitor cell (Lin- CD34+ CD38+ CD45RA- CD123+); GMP: Granulocyte monocyte progenitors (Lin- CD34+ CD38+ CD45RA+ CD123+); MEP: Megakaryocyte-erythroid progenitor cell (Lin- CD34+ CD38+ CD45RA- CD123-); early_PM: Early Promyelocyte (Lin- FSChi SSCint CD34- CD15int CD49dhi CD33hi CD11b- CD16-); late_PM: Late Promyelocyte (Lin- FSChi SSChi CD34- CD15hi CD49dhi CD33hi CD11b- CD16-); MY: Myelocyte (Lin- FSChi SSChi CD34- CD15hi CD49dhi CD33hi CD11bhi CD16-); MM: Metamyelocytes (Lin- FSChi SSChi CD34- CD15hi CD49d- CD33- CD11bhi CD16-); BC: Band cell (Lin- FSChi SSChi CD34- CD15hi CD49d- CD33- CD11bhi CD16int); PMN: Polymorphonuclear cell (Lin- FSChi SSChi CD34- CD15hi CD49d- CD33- CD11bhi CD16hi); Mono: Monocyte (CD14+ CD16-). **B.** Graph represents mean FPKM values for Dmd transcripts encoding Dp427 (red) and Dp71 (blue). While the expression of Dp427 transcript is virtually absent, Dp71 is present in HSC and MPP1, and a significant downregulation of expression with differentiation into MPP3 and MPP4 cells is noticeable; n = 3, # ANOVA p < 0.001, * p < 0.001 vs MPP1.

Therefore, we performed in-depth discovery proteomics in BMM from wild type (WT) and dystrophin-null mdx^βgeo^ mice. To specifically analyse the cell-autonomous alterations, we used BMM on day 7 of differentiation *in vitro.* Using this approach, we compared naïve dystrophic and WT macrophages that were not exposed to any inflammatory signals present in the dystrophic mouse due to the chronic muscle inflammation. Such signals could confound the impact of the *Dmd* gene ablation on macrophages, because of the influence of innate immune memory effect of “priming” these cells on their subsequent reactivity^50,51^.

#### Proteome alterations in dystrophin-null BMM

The protein content was isolated from WT and mdx^βgeo^ BMM and subjected to quantitative bottom-up proteomics using isobaric stable isotope labelling, orthogonal ultra-high performance liquid chromatography, and high-resolution mass spectrometry (iTRAQ 2DLC-MS). The mass spectrometry proteomics data have been deposited to the ProteomeXchange Consortium via the PRIDE [1] partner repository with the dataset identifier *PXD028671*.

We detected 9266 proteins (Supplementary Table 1, Figures 3 A and B) and among the most highly expressed proteins were Grk1 and Pnp, while Pls3 was the most downregulated protein. Overall, a total of 307 proteins were significantly altered between WT and mdx^βgeo^ BMM (FDR corrected *p<0.05*). Post-translationally modified proteins were also profiled from the differential expression of 360 phosphorylated, 1,800 methylated and 500 acetylated proteotypic peptides between the WT and mdx^βgeo^ BMM conditions.

**Figure 3.**
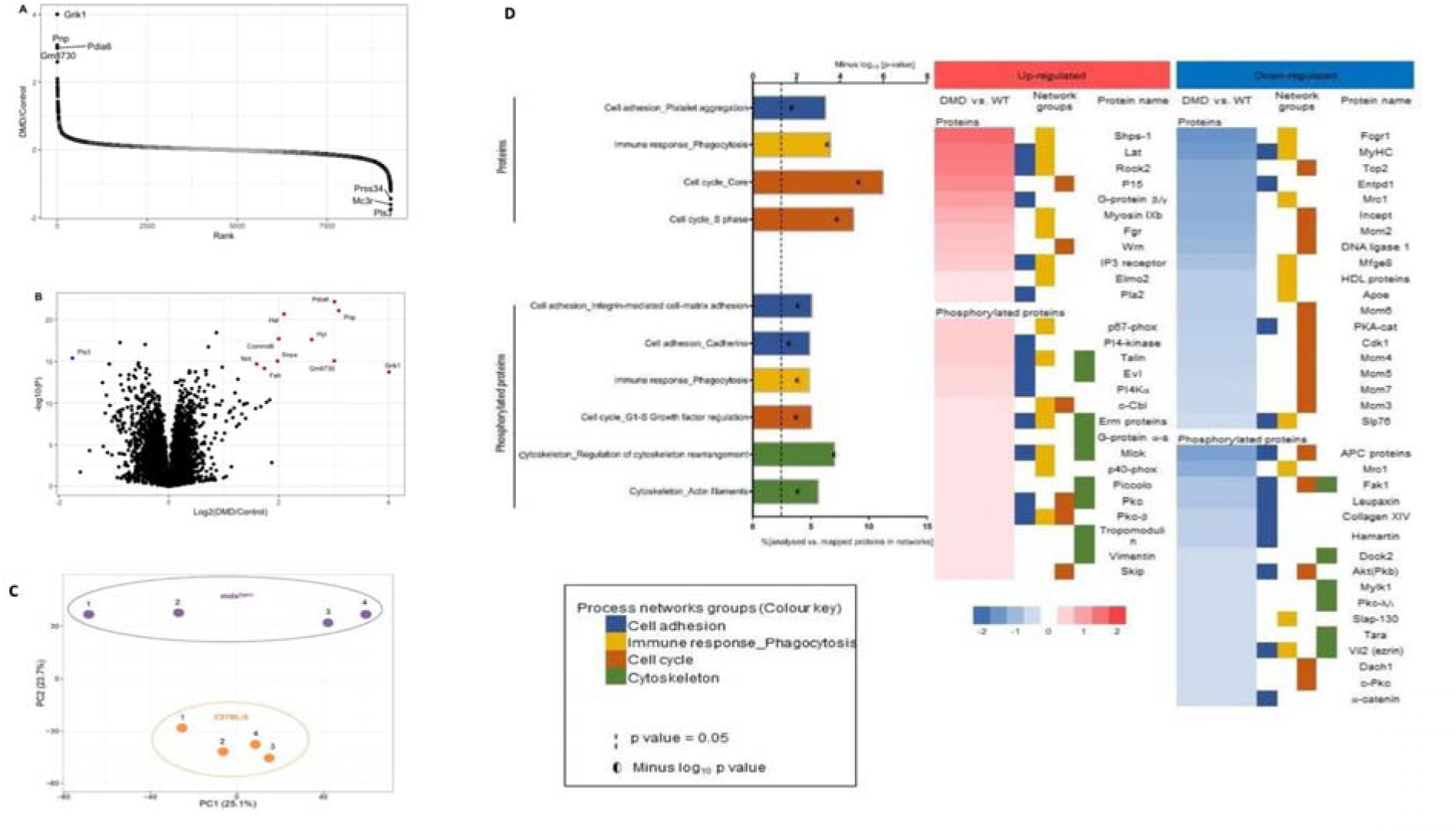
Proteomics analysis of BMM. **(A)** Rank plot displaying protein expression differences between DMD and control mice. Black dots represent significantly expressed proteins, while grey dots represent not-significantly expressed proteins. **(B)** Volcano plot depicting MS results, with proteins coloured in red having a Log2 DMD/Control ratio of > 1.5 and -log10(P) of >10, and proteins coloured in blue having a Log2 DMD/Control ratio of < −1.5 and -log10(P) of >10. **(C)** Principal component analysis using the reporter ion log_2_ratios of all analysed proteins showing mdx^βgeo^ BMM having a distinct proteomic profile and higher heterogeneity compared to C57BL/6 BMMs. **(D)** MetaCore process network analysis showing that process networks related to cell adhesion, phagocytosis, cell cycle and cytoskeleton were over-represented in mdx^βgeo^ BMM.

Principal component analysis (PCA) against the entire proteome profiled showed a clear separation between WT and mdx^βgeo^ BMM mice (Figure 3C). The direct interaction algorithm in MetaCore was used to map altered native proteins and phosphorylated proteins that significantly enrich pathways and their resulting direct protein interaction networks (PIN). Hierarchical clustering of all the significantly altered proteins and their modifications are represented as heatmaps in Supplementary Figure 2 and 3. Functional biological process analysis of differentially expressed proteins showed cell adhesion, phagocytosis and cell cycle networks to be significantly altered in mdx^βgeo^ compared to WT BMM (Figure 3D). The process network analysis of the differentially expressed phosphoproteins, along with the acetylated and methylated proteins, showed consistent enrichment for the cytoskeleton, adhesion, phagocytosis and cell cycle networks (Figure 3D; Supplementary Figures 3 and 4). Specific differentially expressed proteins present in their phosphorylated, methylated and acetylated form were also found overlapping (Supplementary Figure 5).

Analysis of specific proteins within these pathways revealed Plastin 3 (T-Plastin), with a critical role in cell migration^52^ to be the most downregulated protein (log2 FC −1.67, q value=0.00) and P15 protein (cyclin-dependent kinase 4 inhibitor B) as the most upregulated in mdx^βgeo^ BMM (log2 FC 1.16, q value=0.01). However, except for CDK1 significantly downregulated in mdx^βgeo^ BMM (log2 FC −0.39, q value=0.04), other cyclins and cyclin-dependent kinases were either not found or were not significantly differentially expressed. Interestingly, all six minichromosome maintenance proteins (MCM2-7) controlling the DNA replication licensing^53,54^, were found downregulated in mdx^βgeo^ BMM.

The most altered proteins involved in both phagocytosis and adhesion were: Linker of activated T-cells (LAT) (log2FC 1.35, q value=0.00), Rho-associated coiled-coil kinase 2 (ROCK2) (log2FC 1.28, q value=0.00), inositol 1,4,5-trisphosphate (IP_3_) receptor (log2FC 0.43, q value=0.01), and lymphocyte cytosolic protein 2 (Slp76) (log2FC −0.31 q value=0.05).

In addition to Plastin 3, the focal adhesion kinase (FAK), a phosphoprotein involved in cytoskeleton, adhesion and cell cycle pathways, was found significantly downregulated in mdx^βgeo^ BMM. FAK co-localises with talin^55^, and both phosphorylated and methylated talin levels were significantly upregulated in mdx^βgeo^. Other phosphoproteins found significantly upregulated in mdx^βgeo^ were the ERM proteins (ezrin, radixin and moesin) involved in cytoskeleton, cell adhesion and phagocytosis, and the APC protein^56^.

Comparison of proteomic profiles of macrophage markers did not reveal a clear difference in polarisation patterns. The GM-CSF receptor, known to be expressed predominantly on the M1 pro-inflammatory cells, was significantly upregulated in mdx^βgeo^ BMM (log2FC 0.38, q value=0.02). Two M2 macrophage markers, FCγRI and macrophage mannose receptor 1 (CD206), were downregulated in mdx^βgeo^ BMM (log2FC −0.91, q value=0.00 and −0.69 q value=0.00, respectively) and several other markers were not differentially expressed between the two genotypes (Supplementary Figure 6). Importantly, FCγRI and CD206 proteins play major roles in phagocytosis and their downregulation coincided with levels of opsonins and their receptors also being significantly reduced in mdx^βgeo^ BMM (Supplementary Figure 6).

These data suggested an altered migratory and phagocytic activity in dystrophic BMM.

#### Dystrophic bone marrow derived macrophages show key functional abnormalities

Macrophage induced phagocytosis was examined. Fc mediated phagocytosis of fluorescently (CFSE) labelled sheep red blood cells (SRBCs) revealed significant differences between mdx^βgeo^ and WT BMM, with a differential response and different amount of low, medium and highly phagocytic macrophages (Figure 4a), demonstrating impaired phagocytic capability of dystrophic macrophages. Moreover, when stimulated with LPS, significantly fewer mdx^βgeo^ BMM internalised dextran (mol wt 40.000) particles. Whereas in the WT cells dextran uptake gradually increased with the incubation time, in mdx^βgeo^ BMM there was no further increase between 2h and 4h (Figure 4b). Thus, dystrophic macrophages show impairment of phagocytosis and endocytosis.

**Figure 4.**
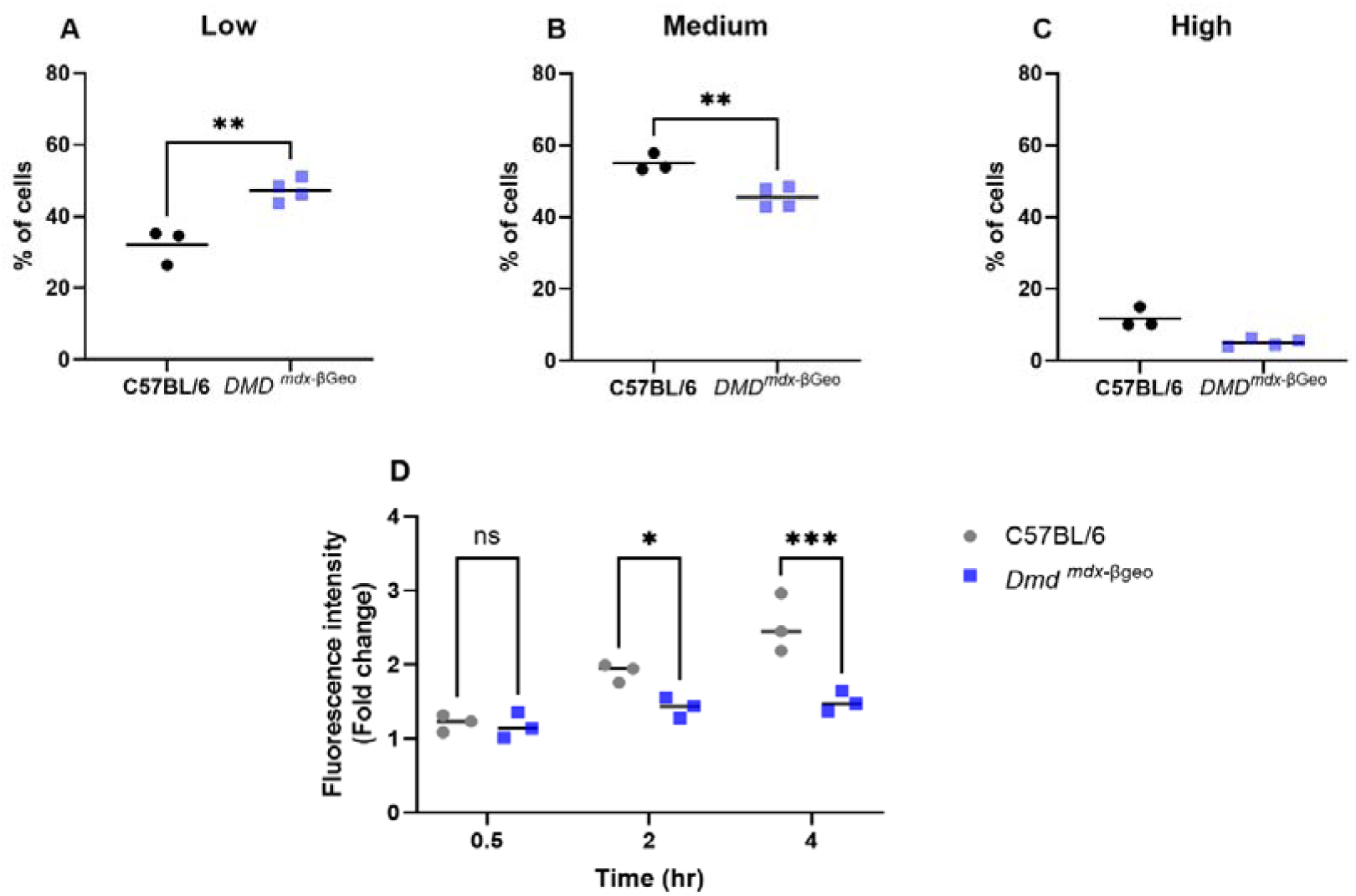
Functional assays of endocytic activity of mdx^βgeo^ BMM. **(A-C)** Fc mediated phagocytosis of IgG-opsonized sheep red blood cells (SRBCs) by BMM. Percentage (%) of cells that were gated as (A) non-/low, (B) medium or (C) highly phagocytic are shown on the graphs as single data points, mean ± SEM (unpaired t-test). **(D)** Endocytosis of dextran-FITC (40.000 MW) particles by LPS-stimulated BMM. Fluorescence intensity (corresponding to dextran uptake, values normalized to control C57BL/6 BMM at 0.5h in each experiment) in cells, following 0.5-, 2-, and 4-hour incubation with dextran, is shown on the graphs as single data points, mean (2way ANOVA with Šídák’s multiple comparisons test). *p <0.05

The altered Process Networks Groups for cytoskeletal proteins were another key finding in the proteome of mdx^βgeo^ macrophages. Confocal analysis of F-actin and alpha-tubulin cytoskeleton components showed networks of these proteins to be significantly disorganised: The expected F-actin distribution primarily around the cell periphery present in the WT cells was lost in mdx^βgeo^ BMM (Figure 5a). Moreover, the majority of mdx^βgeo^ BMM were flattened, with the average thickness <3 µm being about half of that in WT cells (Figure 5b). These cytoskeletal rearrangements could result in altered cell motility. Indeed, mdx^βgeo^ BMM ability to migrate in the Boyden chamber assay was significantly (2.3-fold) reduced compared to the WT cells (Figure 6c). The mdx^βgeo^ BMM invasiveness in the gelatine-coated chamber also appeared reduced, with 1.6-fold decrease. However, with *p = 0.058* there was not enough power to prove the alternative hypothesis (Figure 5c).

**Figure 5.**
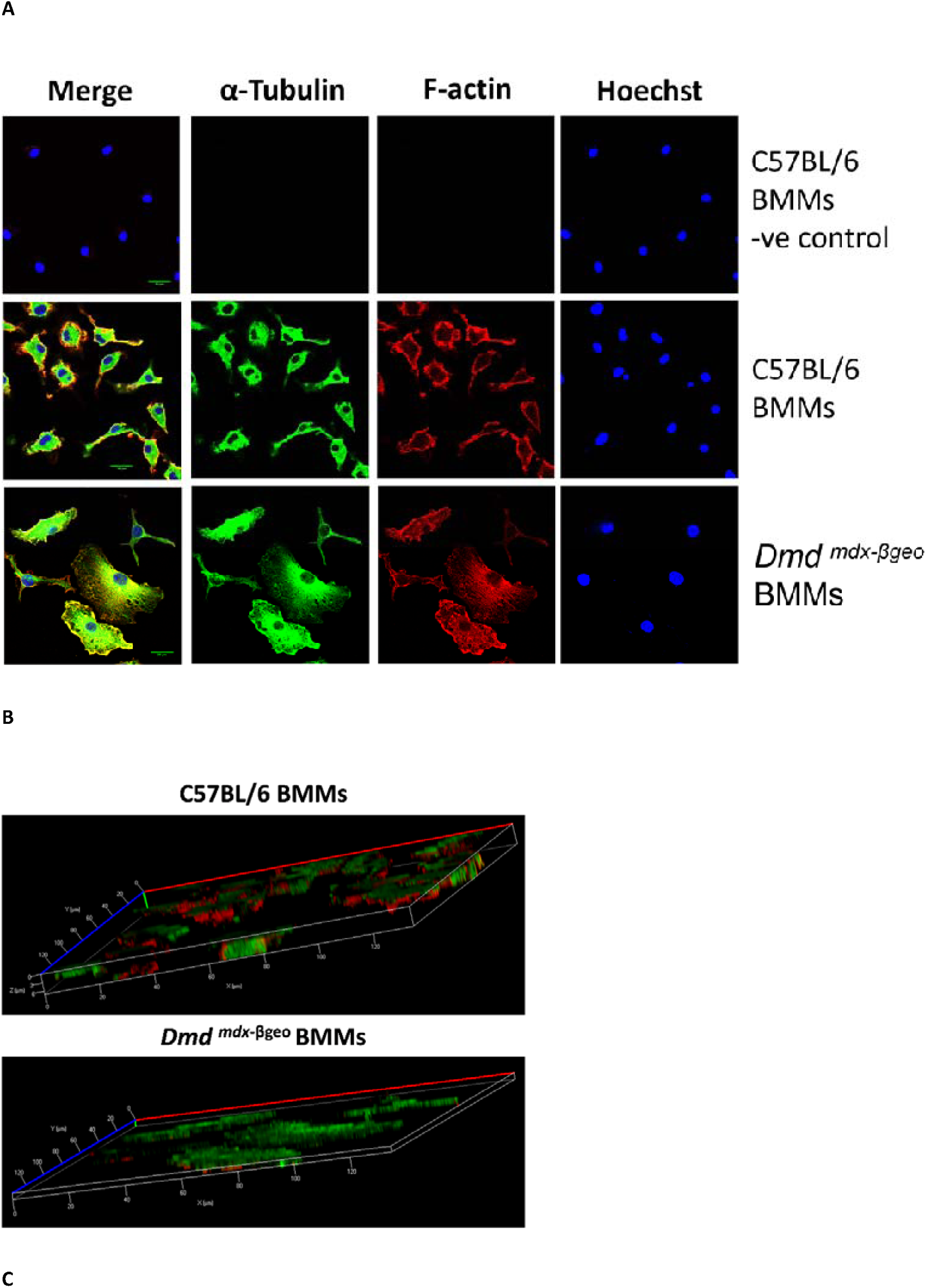
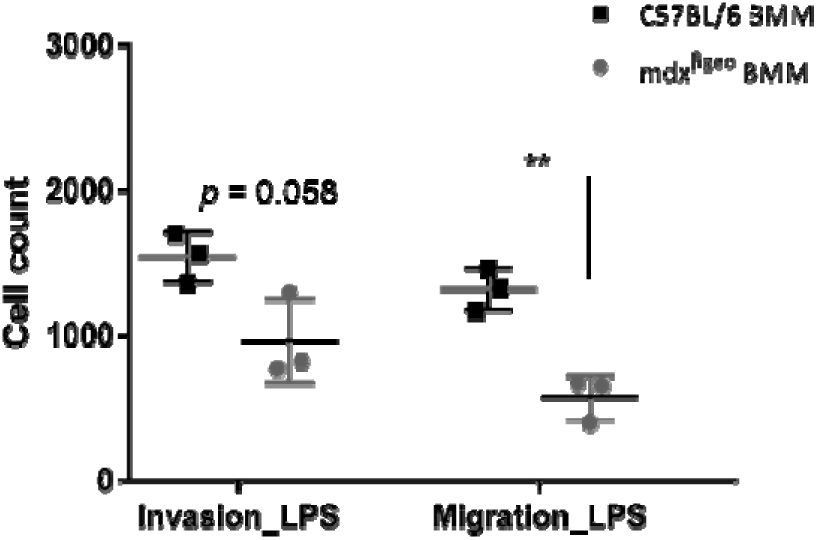
Cytoskeleton alterations in mdx^βgeo^ BMM. (**A**) Examples of confocal images of macrophage cytoskeletons stained for alpha-tubulin (green), F-actin (red) and nuclei stained with Hoechst (blue). In the negative control (-ve), secondary antibody only was used to ensure specificity of the signal. Scale bar = 20µm. **(B)** 3D confocal image rendering of BMM cytoskeletons. The staining is as in A. Thickness (Z) of C57BL/6 cells was around 6 µm while in mdx^βgeo^ cells it was less than 3 µm. **(C)** BMM migration and invasion assays. Averaged numbers of C57BL/6 and mdx^βgeo^ BMMs, which migrated through the gelatine-coated filter (invasion) or uncoated Boyden chamber membrane (migration) are shown. n ≥ 3, mean ± SD, **p = 0.01

**Figure 6:**
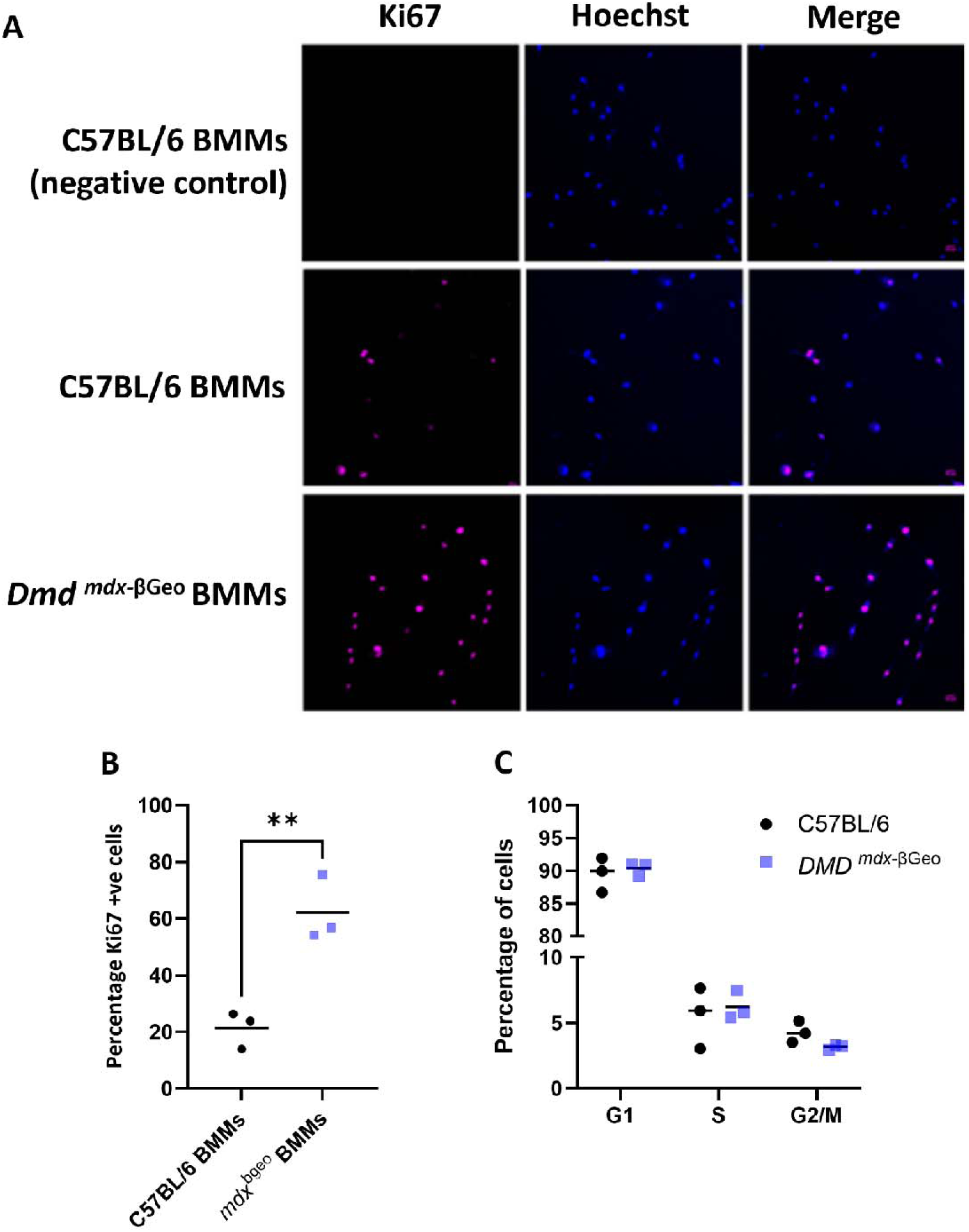
Altered **c**ell proliferation profile in dystrophic BMM. **(A**) Comparative confocal images of Ki67 staining in BMMs: Ki67 immunofluorescence in magenta, cell nuclei stained blue with Hoechst; the negative control (-ve) with secondary antibody only was used to ensure specificity of the signal. (**B)** The percentage of cells from each genotype expressing Ki67; n = 3, **=p < 0.01. **(C) T**he proportion of cells in different phases: G1, S, and G2/M of the cell cycle for C57BL/6 and mdx^βgeo^ BMM.

Although the functional differentiation of cells is known to reduce their proliferative capability, macrophages have some ability to proliferate^57,58^. Notably, the cell division protein networks (Cell Cycle_Core, Cell_Cycle S-Phase) were found significantly altered in mdx^βgeo^ BMM (Figure 3b). Ki67 analysis showed 3-fold increase (p=0.01) in the number of mdx^βgeo^ BMM that expressed this marker of cell proliferation (Figure 6a, b). However, no significant difference was found in the cell cycle progression between mdx^βgeo^ and WT BMM: ∼90% of cells were in G0/G1 phase, only ∼ 5.5% of cells were in S phase and ∼4% of cells were in G2/mitosis (Figure 6c). In the Bromodeoxyuridine (BrdU) incorporation test, detecting DNA synthesis as a measure of cell proliferation, correspondingly small proportion of BMM was found to be labelled and no significant difference between the mdx^βgeo^ dystrophic and WT cells was observed.

In response to tissue damage, macrophages secrete a variety of inflammatory mediators, which activate defence mechanisms directly and engage other cells to sustain the inflammation^59^. The macrophage NLRP3 inflammasome is important for establishing the inflammatory response following activation of the purinergic P2X7 receptor^60^. Interestingly, increased P2X7 expression and function have been found in dystrophic myoblasts and lymphoblasts^27^. Western blotting analysis of protein extracts from fully differentiated and unstimulated BMM revealed that P2X7 and NLRP3 proteins were significantly overexpressed in mdx^βgeo^ compared with WT BMM (Figure 7a and b). While pre-IL-1β expression, as expected, showed a dose-dependent increase in LPS-activated macrophages, levels of this interleukin were significantly higher in mdx^βgeo^ compared to WT cells (Figure 7c).

**Figure 7.**
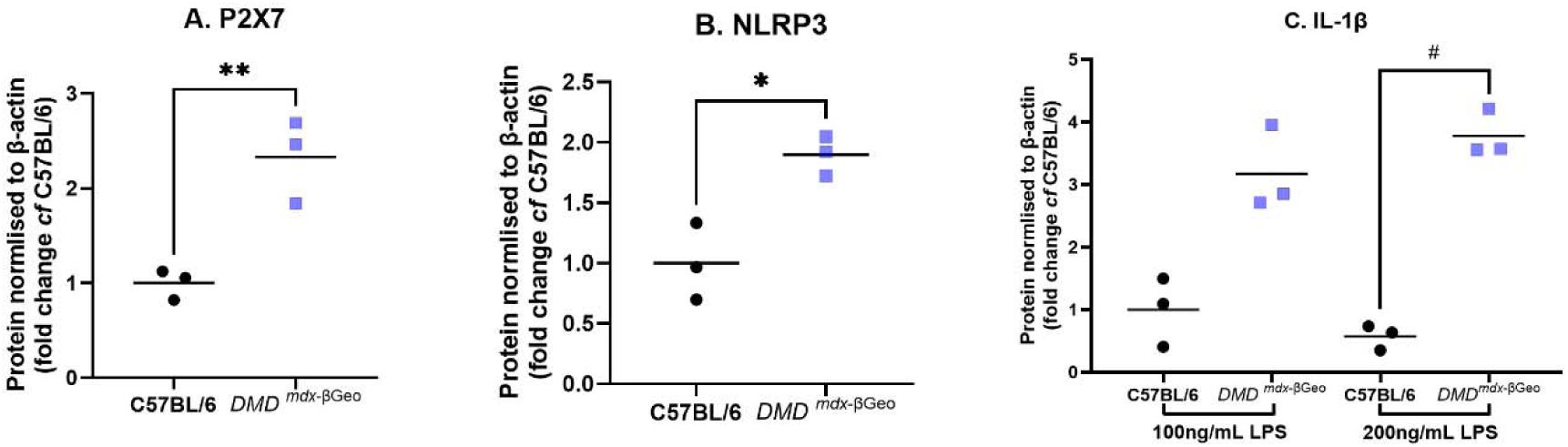
Inflammatory mediator alterations in mdx^βgeo^ BMM. Densitometric analysis of western blot data revealing significantly higher protein expression levels for (A) P2X7 (** = p<0.01, t-test), and (B) NLRP3 (*=p<0.05) in BMM. (C) Expression of pro-IL-1β increasing significantly in mdxβgeo BMM stimulated with rising amounts of LPS (# = p<0.05, Kruskal-Wallis test with Dunn’s post-tests).

In contrast to the protein data, RT-qPCR analysis of IL-1β and NLRP3 transcripts, showed great diversity of expression of these genes in mdx^βgeo^ but no significant difference in expressions in mdx^βgeo^ compared to WT BMM (not shown). This may suggest a post-transcriptional or epigenetic mechanism of alteration.

Notably, the majority of DMD patients has mutations upstream from Dp71, which therefore should not affect BMM functions. Interestingly, RT-qPCR analysis in BMM derived from the mdx mouse, which has the stop codon mutation in exon 23 thus disrupting full-length dystrophin expression, showed a statistically significant upregulation of Dp71 transcript levels in early developing BMM (Figure 8a). Given that overexpression of Dp71 has been found to worsen the disease phenotype^61,62^, we analysed phagocytosis /endocytosis in mdx BMM. However, these key functional alterations found in mdx^βgeo^ cells were not observed in mdx macrophages (Figure 8b, c and Supplementary Figure 7).

**Figure 8.**
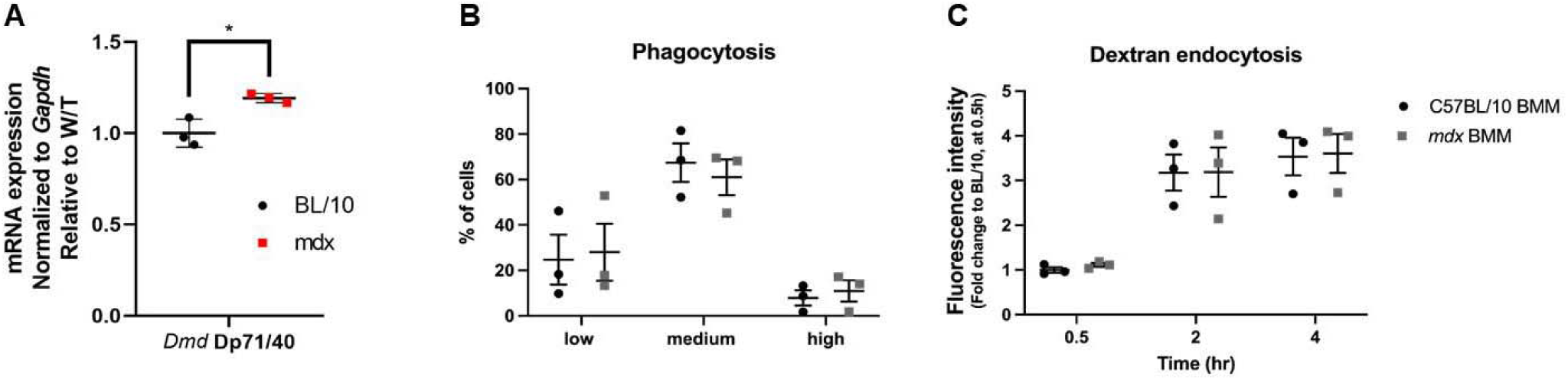
The mdx macrophages are not functionally altered. **(A)** Increased Dp71 expression in differentiating mdx BMM, n=3, *p < 0.05. **(B)** Fc mediated phagocytosis of IgG-opsonized sheep red blood cells (SRBCs) by BMM. Percentage (%) of cells that were gated as non-/low, medium or highly phagocytic are shown as single data points. **(C)** Endocytosis of dextran-FITC (40.000 MW) particles by LPS-stimulated BMM. Fluorescence intensity (corresponding to dextran uptake, values normalized to WT C57BL/6 BMM at 0.5h in each experiment) in cells, following 0.5-, 2-, and 4-hour incubation with dextran, is shown as single data points. For B and C: mean ± SEM (unpaired t-test with Welch’s correlation) shown.

#### Loss of DMD gene expression affects peritoneal macrophages

Macrophages residing in most adult tissues appear during the embryonic development, have several origins in ontogeny but not originate (at steady state) from circulating monocytes^63^. Moreover, these cells are highly plastic and exhibit great functional diversity. To assess whether loss of DMD gene expression impacts both BMM and tissue-resident macrophages, we compared peritoneal macrophages (PMϕ) from two dystrophic and their respective WT strains. Given that ability to adhere and migrate into and out of a tissue is key to macrophage function, and that we identified a migration deficit in mdx^βgeo^ BMM, we used live cell imaging to assess motility of these cells. Motility was found to be significantly impaired in dystrophin-null PMϕ when compared to WT (Figure 9), with many mdx^βgeo^ cells showing poor attachment (Figure 9a) and no significant movement over the entire 24h measurement period (Figure 9b-d). In contrast, the motility of mdx PMϕ was not affected (Figure 9), in agreement with BMM results.

**Figure 9.**
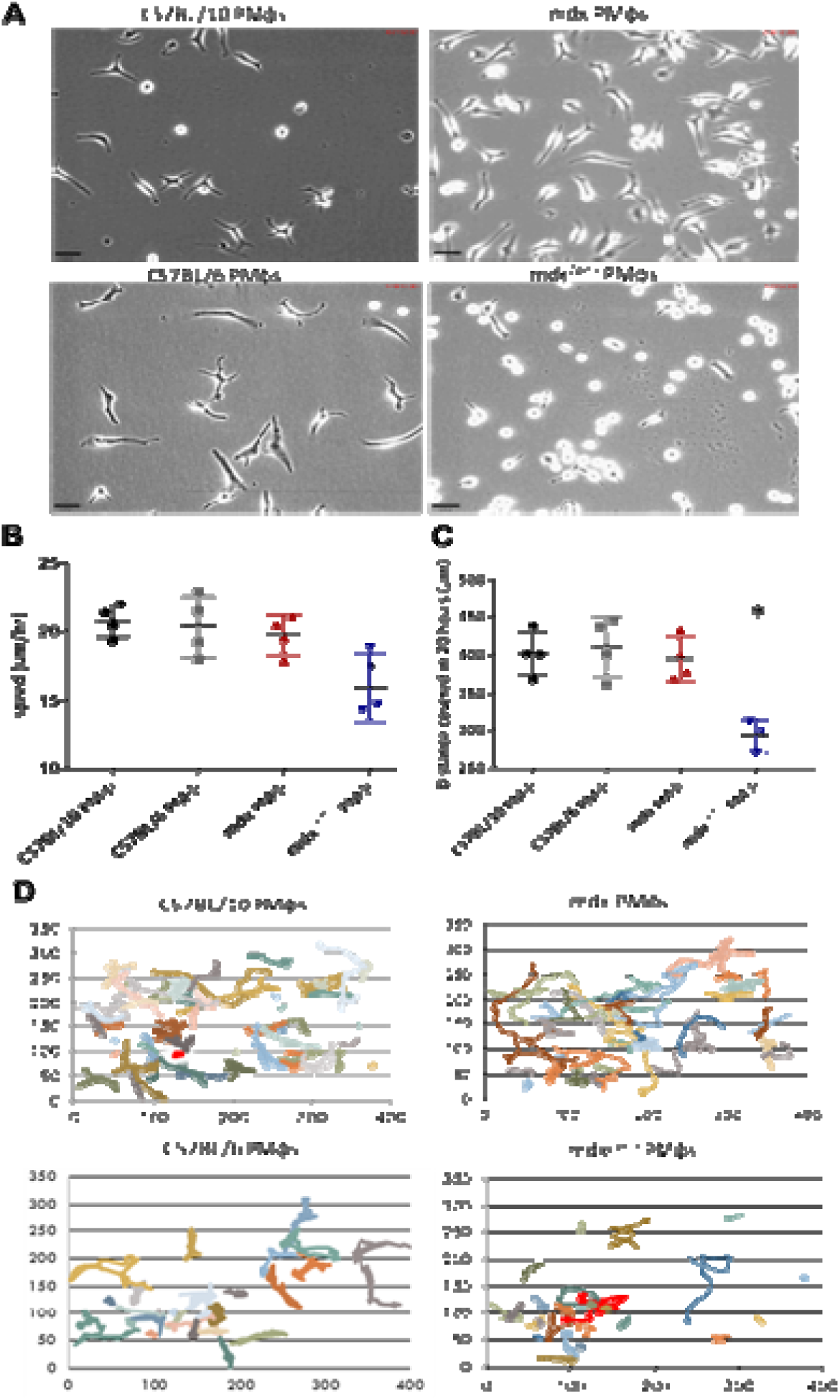
Cell adhesion and migration of PMϕs. (**A**) Bright field microscopy images of cells after 20 hours in culture. An abnormally rounded cell body with the lack of the protrusions is clearly noticeable in mdx^βgeo^ PMϕs. Scale bar = 30 µm. Analysis of migration of PMϕs using live cell imaging. **(B)** The average distance covered by wild-types (C57BL/6 and C57BL/10) and dystrophic (mdx and mdx^βgeo^) PMϕs in 20 hours. **(C)** Individual value plots illustrating the speed of WT and dystrophic PMϕs. For A and B n ≥3, mean ± SD, *p <0.05. **(D)** Example diagrams showing the migration paths (μm) of individual macrophages.

In summary, Dp71 dystrophin plays an important role in the development and function of mouse macrophages. Although Dp71 absence does not preclude their differentiation, dystrophic BMM present with key functional alterations. Dystrophin-null PMϕ are also functionally altered.

## DISCUSSION

We describe here, for the first time, cell-autonomous abnormalities in dystrophic macrophages resulting from the loss of expression of specific dystrophin transcripts. Healthy HSC and developing bone marrow macrophages expressed Dp71 mRNA, but the levels of expression decreased with cell differentiation into mature BMM. Loss of *Dmd* gene expression in mdx^βgeo^ manifested in significant proteome alterations in BMM and was detrimental to key Mϕ functions, such as migration, phagocytosis and the NLRP3 inflammasome.

These defects were cell autonomous: They manifested in naïve BMM differentiated from bone marrow stem cells *in vitro* and therefore never exposed to the inflammatory mediators in the dystrophic muscle. Moreover, *mdx* macrophages, which express Dp71 and have been exposed to the dystrophic inflammatory environment, were not altered in a similar manner. Lack of abnormalities in *mdx* cells, where a *stop* mutation eliminates only full-length dystrophins, confirms the significance of Dp71 short isoform in macrophages.

A notable proteomic alteration in mdx^βgeo^ BMM, was Plastin 3 (T-Plastin) that was found to be the most downregulated protein within the altered pathways. Plastin 3 cross-links actin microfilaments that support cell shape, movement and plasticity^52^. These traits were in line with functional alterations observed in dystrophic macrophages.

Significant downregulation of the Minichromosome Maintenance (MCM) proteins in dystrophic macrophages is also notable given that even a mild reduction in the MCM pool compromises DNA replication^64^ and results in genome instability^65^. It also impacts cell migration^66^ and reports suggested a critical role for these proteins in NK cells and the NK cell–mediated host defence^67^. Of note, MCM expression has been associated with increased Ki67 proliferation marker^54^, while we found MCM downregulation coinciding with a 3-fold increase in Ki67 expression but no obvious cell proliferation alterations in dystrophic macrophages. While Ki-67 is a marker of proliferating cells and is absent in completely quiescent or senescent cells, it is detectable at low levels in recently quiescent cells^68^. Therefore, the higher Ki-67 may suggest that mdx^βgeo^ exited the cell cycle later than WT BMM. Further studies should shed some light on the significance of these findings.

Importantly, proteomic results predicted altered migration, phagocytosis and the NLRP3 inflammasome, and functionally confirmed. While the focus of DMD research has, understandably, been on muscle cells, macrophages are known to be involved in this disease progression. Inflammation has been shown to precede muscle damage in DMD patients^23–25^ and early macrophage depletion reduced the dystrophic phenotype^69^. Yet, while their targeting alleviated symptoms^70^, total ablation exacerbated the disease^69^, in agreement with the role these cells play in muscle regeneration^71^. Skeletal muscle injury activates resident macrophages and recruits monocytes. Initially displaying an inflammatory profile, upon phagocytosis of muscle cell debris, Mϕ switch to an anti-inflammatory phenotype that stimulates muscle regeneration^72^. Therefore, abnormal migration and phagocytosis found in dystrophin-null Mϕ could affect their ability to remove the muscle debris. Moreover, as phagocytosis induces a switch toward an anti-inflammatory phenotype^72^, a phagocytic defect could perpetuate the inflammatory status. Cell surface marker profiles in mdx^βgeo^ BMM were confusing, with some normally associated with anti-inflammatory phenotype contrasted by significantly upregulated GM-CSF receptor levels, expressed predominantly on M1 cells. Altered macrophage phenotype transition can impair regeneration of skeletal muscles. This, combined with overexpressed NLRP3 inflammasome components in dystrophic Mϕ (Figure 7), could further potentiate muscle inflammation, with TNFα released by macrophages triggering necroptosis in dystrophic muscle cells^73^.

However, it must be stressed that this autonomous abnormality in dystrophin-null macrophages is not a key factor in DMD pathogenesis. 80% of mutations in DMD patients occur in two hotspots disrupting the full-length dystrophins only^74^ and no analogous abnormalities were found in *mdx* macrophages. Thus, the loss of full-length dystrophins is both necessary and sufficient for DMD to occur. Yet, the dystrophin-null mutations impacting both muscle and non-muscle cells, including macrophages, could lead to the most severe phenotype^38,75^. Indeed, increased ectopic calcification and fibrosis associated with altered macrophage infiltration patterns was found in mdx^βgeo^ ^39^ and macrophages have been implicated as direct contributors to fibrosis^76^.

Interestingly, while proper myogenic cell functions depend on the expression of full-length dystrophin transcripts and macrophage functions show Dp71 dependency, loss of expression of these vastly different dystrophins leads to some common cell-autonomous abnormalities in these two cell types: Proteomic analyses in Mϕ (this work) and myoblast^77^ identified alterations in the ERM (ezrin, radixin and moesin) proteins and their functionally-relevant phosphorylation, which changes were linked to altered migration of these cells. Furthermore, both cell types manifest the P2X7 purinergic phenotype^78^. Interestingly however, while Dp427 loss in all cells displaying a phenotype is associated with increased intracellular calcium levels^79^, we found no differences in store operated calcium entry between mdx^βgeo^ and WT macrophages (Supplementary Figure 8). On the other hand, loss of Dp71f in tumour cells affected calcium influx^10^. This suggests that Dp71 functions may be isoform-dependent and/or involve calcium homeostasis in cell-specific manner.

Given that Dp71 expression showed significant downregulation as HSC differentiated towards myeloid and lymphoid cells (Figure 2b, Suppl. Figure 1), we investigated whether the opposite happens in malignant cells, which retain a degree of undifferentiated phenotype. Indeed, analysis across two databases revealed statistically significantly increased overall expression of Dp71 and of its exon 78 splice variant (Dp71b) across haematological malignancies compared to normal blood (Supplementary Figure 9). This further links Dp71 with the undifferentiated phenotype and supports its significance there.

However, the mechanism in which loss of *Dmd* gene expression leads to abnormalities in macrophages is puzzling. While we detected Dp71 transcript in wild type BMM and PMϕ, its levels in mature cells were low and no dystrophin could be detected by Western blotting. Yet, dystrophic macrophages manifested clear abnormalities. One possible explanation is that dystrophin is expressed in bursts at specific checkpoints. Small burst of expression would not be easily detectable in non-synchronised macrophages in culture.

Yet, significantly higher *Dmd* expression in HSC and early differentiating hematopoietic cells than in macrophages may suggest another explanation, which involves alterations of epigenetic regulation. This mechanism is of increasing importance in DMD, where loss of gene expression in muscle stem cells affects not only their functions^80–82^ but the descendant myoblasts harbour somatically heritable epigenetic changes^15^ and manifest significant epigenotype abnormalities^83^. Moreover, in dystrophic macrophages, the phenotypic changes might be the consequence of some events that occurred in their development and then become irreversible after crossing a specific checkpoint. The overall similarity between the *Dmd* dysfunction in muscle stem cells affecting myoblasts and in HSC and macrophages suggests that mis-regulation in progenitors might be the common outcome of *DMD* gene mutations across tissues, despite involving two different isoforms: Dp427 and Dp71.

Another, related, possibility is that loss of dystrophin expression triggers a pathological mechanism affecting RNA rather than its protein product. Mutant dystrophin mRNA degradation is known to be inefficient, especially for the distal mutations. It has been reported that mutations downstream of exon 70 can evade degradation^84^. In fact, recent studies revealed that premature termination codons affect DMD transcripts not, as expected, *via* nonsense mediated decay (NMD) but through a possible epigenetic mechanism resulting in a chromatin conformation that is less open for transcription. The mutant dystrophin mRNA localizes in cell nuclei. These data suggest that the mutant DMD locus causes regional changes in transcriptional dynamics likely mediated by histone marks^85^. Such mutant RNA might be altering transcriptional control of other protein-coding as well as non-coding genes (including lncRNAs and miRNAs). Moreover, the DMD locus encodes specific lncRNAs^86^ and DMD deletion could deregulate expression of such lncRNAs, which, in turn, can trigger additional abnormalities e.g. by altering miRNA sponging^86–88^. In fact, in early myogenic cells, significant miRNA alterations were found^15^.

Interestingly, human lymphoblasts, which when healthy express *DMD* transcripts but no detectable protein, were found to exhibit purinergic abnormalities, when dystrophic^22^. The same purinergic phenotype was found in dystrophic myoblasts^78,89^.

These multiple yet diverse transcriptomic alterations in dystrophic myoblasts, myotubes, lymphoblasts and now in macrophages could be explain by transcriptional adaptation, a recently discovered cellular response, that has been just identified in DMD^90^. In this process, independent of the loss of protein function, mutant mRNA decay leads to the increased transcription of “adapting genes”^91^.

In macrophages, an abnormality due to the lack of dystrophin expression in the stem cell continuing downstream, aligns with the Dp71 developmental expression pattern overall: it is the first product of the *DMD* gene associated with morphogenic events and terminal differentiation of various mouse cells and organs^41,45–47,92^.

Altogether, our data expand the growing body of findings that the DMD pathology is a progressive developmental disease^15^, with an embryonic onset^15^. Functional abnormalities appear to affect development of muscle and non-muscle stem cells and persist in their progeny. Importantly, consequences of *DMD* mutations are truly widespread. Apart from muscle, they also cause defects in the development of neurons and glia as well as cells derived from bone marrow progenitors, such as macrophages (this work), lymphocytes^22^, megakaryocytes (manifesting in altered aggregation of dystrophic platelets)^6^ and also endothelial cell abnormalities^93^. We recently found dystrophin to be expressed across human tissues and DMD downregulation occurring in tumours coinciding with Duchenne-like molecular alterations^8^. We postulate that epigenetic dysregulation and/or transcriptional adaptation in dystrophic stem cells triggers a pathological cascade with the functional consequences observed in daughter cells.

In humans, this developmental dysfunction appears linked to the loss of expression of the specific embryonic dystrophin^17^, but in lower species, not expressing Dp412e, one of the other isoforms could be involved. It is possible that this *DMD* gene divergence could contribute to the species variability in disease phenotypes. Likewise, differences between human and mouse innate immunity and specifically in macrophages^94–96^ need to be investigated to fully understand the impact of the loss of DMD gene expression in these cells. However, it is important to remember that both human and mouse HSC and macrophages express Dp71.

In conclusion, the normal development and function of macrophages requires Dp71 expression. Early absence of this dystrophin alters migration, phagocytosis and the NLRP3 inflammasome, which are key Mϕ functions intricately involved in the dystrophic process. Therefore, comparisons of dystrophin-null patients and those suffering from mutations affecting full-length dystrophins are warranted, as these pathological phenotypes may differ and benefit from a modified treatment.

## MATERIALS AND METHODS

### Animals

Male C57BL/10ScSn-*Dmd^mdx^*/J, C57BL/10ScSnJ, C57BL/6-DmdGt(ROSAbgeo)1Mpd/J (*Dmd^mdx-^*^βgeo^) and C57BL/6J eight-week-old mice were used in accordance with institutional Ethical Review Board and the Home Office (UK) Approvals. All mice were maintained under pathogen-free conditions and in a controlled environment (12-hour light/dark cycle, 19-23°C ambient temperature, 45-65 % humidity). For environmental enrichment, tubes, toys and nesting materials were routinely placed in cages.

Dmd^m*dx-βge*o^ mice were generated in C57BL/6 by insertion of the ROSAβgeo promoter-less gene trap construct downstream from the dystrophin DP71 promoter, resulting in the reading-frame disruption and loss of all dystrophin isoforms^40^ and P2rX7^-/-^ knockout mice were as described^97^. The C57BL/10 and C57BL/6 strains derived from a common origin^98^ and it has been demonstrated that the mdx mutation on the C57BL/6 background shows the same pathology as the original C57BL/10 strain^99^. The genotypes of experimental animals were confirmed by PCR (Supplementary Figure 10).

### Macrophage isolation

#### Isolation of peritoneal macrophages

Mice were killed by CO_2_ exposure, abdomens sterilised with 70% ethanol and 5 ml cold DPBS was injected into the peritoneal cavity. Peritoneum was then gently massaged, and fluid was aspirated. Collected peritoneal cells were centrifuged at 300 g for 8 min at 4°C. Supernatant was discarded and pelleted cells were re-suspended in 3ml DMEM F12 with 10% (v/v) FBS, 2mM L- Glutamine, 100U/ml Penicillin and 100U/ml Streptomycin (Fisher Scientific, UK). Cell suspension was plated in polystyrene petri dish and incubated at 37°C in a humidified environment of 5% CO_2_ for 20 min or O/N. Non-adherent cells were then discarded.

#### Isolation of bone-marrow derived macrophages

Femur and tibia were dissected and cleaned mechanically using a paper tissue to remove the surrounding muscles. Bones were placed in cold DPBS in a petri dish and transferred to a sterile hood. After soaking for 30 sec in 70% ethanol and two washes in sterile PBS, the contents of bone marrow cavities were flushed out with cold PBS using a syringe and a needle. Eluates were passed through a 70 µm cell strainer into falcon tubes and centrifuged at 300 g for 8 min at 4°C. Supernatant was discarded and cell pellet was re-suspended in 3ml ACK lysis buffer (Life technologies Ltd.) and incubated for 5-10 min at RT for red blood cell elimination. 3ml PBS was added to stop the reaction and cells were centrifuged at 300 g for 6 min. Supernatant was discarded and pellet re-suspended in 10ml DMEM F12 with 10% (v/v) FBS, 2mM L- Glutamine, 100U/ml Penicillin, 100U/ml Streptomycin and 20ng/ml Macrophages-Colony Stimulating Factor (M-CSF) (Sigma Aldrich Ltd.). Suspension was plated in 10 polystyrene petri dishes or tissue culture 6-well plates (Fisher Scientific Ltd.) and incubated at 37°C in a humidified environment of 5% CO_2_. On day 3, 3 ml of the same fresh medium was added to each petri dish. On day 7, non-adherent cells were removed, and adherent cells were considered macrophages.

### RNA extraction, cDNA synthesis and qPCR analysis

Total RNA was extracted from *Dmd*^mdx^, *Dmd*^mdx-βgeo^ and control cells using RNEasy Plus Universal mini kits (Qiagen 73404). Cells in the culture dish were washed 2 times with warm PBS (Gibco 14190144) before addition of the lysis buffer and suspension passed through a 25-gauge needle for 20 times. Samples were then processed according to kit manufacturer’s instructions. RNA quality and concentration were measured using a NanoDrop 1000 Spectrophotometer (Thermo Scientific). RNA integrity was assessed using electrophoresis of 100 ng of total RNA in a 1 % agarose gel (Sigma A4718) in TAE buffer or using an automated capillary electrophoresis system (2100 Bioanalyzer Instrument G2939BA, Agilent) using a kit assay (Agilent RNA 6000 Nano Kit 5067-1511, Agilent).

Total RNA samples were converted to cDNA using SuperScript VILO cDNA Synthesis Kit (Invitrogen 11754050) as per manufacturer instructions.

#### Reverse Transcriptase-Polymerase Chain Reaction

25-100 ng of each cDNA were included in a 25µl reaction volume containing 200-500nM of specific primers (Supplementary Table 2), with 2.5 units of *Taq* DNA Polymerase (*Taq* DNA Polymerase kit from Qiagen Ltd.) in 1X CoralLoad PCR Buffer. Samples were amplified using either Primus 25 42 Advanced ® or Primus 96 Plus® (PEQLAB Biotechnology GmbH, Erlangen, Germany) with PCR cycling profile: 94°C/ 3 min, followed by 35 cycles of 94°C/30-60 sec, appropriate annealing temperature for 60 sec (Supplementary Table 2) and 72°C/60 sec/kb of product size, with a final extension step of 72°C for 10 min. PCR products (10µl) were resolved on 2% agarose gels and visualised using the Gel Doc^TM^ EZ Imager (Bio-Rad Ltd.).

#### Real Time Quantitative PCR (RT-qPCR)

SYBR Green qPCR reactions were ran in duplicates using 25 ng of cDNA per reaction with Precision Plus Mastermix (Primer Design PPLUS-LR), forward and reverse primers (Supplementary Table 2) (Eurofins) and DEPC treated water (Fisher Bioreagents BP561) as per manufacturer instructions, on 96 well plates using an Applied Biosystem ViiA7 RT-PCR instrument and expression quantified using the ΔΔCT method. For Taqman: 25ng of cDNA was added to PCR tubes containing a mixture of 10µl of 1x TaqMan Universal Master Mix II with UNG (Applied Biosystems) and 1µl of the 20x specific TaqMan hybridisation probe in the final reaction volume of 20µl. All Taqman primers were purchased from Life Technologies Ltd. (Supplementary Table 2).

### Live Cell Imaging

Cells were seeded at 70-80% confluency in 6 well plates pre-treated with Nunclon™ Delta surface (Sigma Aldrich Ltd.) in 2 ml DMEM F12 with 10% (v/v) FBS, 2mM L- Glutamine, 100U/ml Penicillin and 100U/ml Streptomycin medium and left to attach overnight. The next day, supernatant was discarded, and plates were washed with 2ml warm PBS to remove any non-adherent cells. 2ml of normal medium was again added and plates were immediately placed on a normoxic environment platform of the live cell-imaging microscope (Axiovert 200M; Carl Zeiss Oberkochen, Germany) for 24 hr. To record cell movements, images were taken every 5 min. Statistical analysis (one-way ANOVA with Tukey post-hoc analysis, Minitab 17) was then applied to the series of images of cell positions and movements.

### Immunoblotting

Cells were scraped off in a minimal volume of extraction buffer composed of 1x LysisM extraction buffer (Cat. #04719956001, Roche), 1 protease inhibitor cocktail tablet, 1 phosphatase inhibitor cocktail tablets (both Roche) per 10ml of buffer. Samples were homogenised by passing through a 25-gauge needle 20 times, centrifuged and protein concentrations of extracts determined using the bicinchoninic acid kit (Sigma-Aldrich Ltd). 30-50µg of protein were mixed with Laemmli sample buffer (Bio-Rad Laboratories Ltd.) supplemented with 5% (v/v) β-mercaptoethanol, heated at 95°C for 5 min and chilled on ice before being separated by electrophoresis in polyacrylamide gels. Samples were electrotransferred onto methanol-activated polyvinylidene fluoride membranes (Amersham System, GE Healthcare Life Sciences Ltd.). Membranes were incubated on a horizontal shaker for 1 hour at 37 °C in a blocking solution containing Tris-buffered saline with 0.1% Tween (ThermoFisher Scientific) and 5% dry non-fat cow milk. Membranes were subsequently incubated with primary antibodies diluted in the same blocking solution overnight at 4 °C. Membranes were washed (4 x 5 min) in TBS-Tween and incubated with appropriate HRP-conjugated secondary antibodies in the blocking solution for 2 hours. Finally, membranes were washed 3 x 30 min and incubated with LuminataTM Forte chemiluminescence development reagent (Merck, Burlington, MA, USA) and subsequently imaged using a ChemiDoc MP system (Bio-Rad, Hertfordshire, UK).

### Bioinformatic analysis of DMD transcript expression levels

To analyse mRNA abundance levels of DMD isoforms in HSC cells and MPPs we used raw RNA-seq data available in the ArrayExpress database (http://www.ebi.ac.uk/arrayexpress) under accession number E-MTAB-2262 (Ref^44^). The RNA-seq reads (from FASTQ files) were aligned to *Mus musculus* reference genome (GRCm38 Ensembl version 100) using Hisat2 2.2.1. The transcript FPKM (Fragments Per Kilobase of transcript per Million fragments mapped) levels were quantified and normalized using the Cufflinks v2.2.1 and the modified GTF from the Ensembl gene database. The abundance levels of the transcripts encoding DP71 (ESMUST00000239019) and DP427 (ENSMUST00000114000) were compared.

The UCSC Xena Functional Genomics Browser (http://xena.ucsc.edu) was used to compare the expression of Dp71 and its splice variants between healthy whole blood and primary blood malignancies using the dataset “RSEM expected_count” of the TCGA TARGET GTEx cohort. Four comparisons were performed between whole blood samples (n= 366) from the GTEx database (https://gtexportal.org/) and acute myeloid leukaemia (n= 173) and diffuse large B-cell lymphoma samples (n= 47) generated by the TCGA Research Network (https://www.cancer.gov/tcga) and acute lymphoblastic leukaemia (n= 37) and acute myeloid leukaemia samples (n= 29) generated by the TARGET initiative (https://ocg.cancer.gov/programs/target). Transcript IDs were matched to specific *DMD* transcripts using the Ensembl database. A two-tailed Welch’s t test was performed using the Xena Browser and p-values were adjusted for multiple testing using the Bonferroni correction; they were multiplied by the number of comparisons (n= 4) and compared to the overall level of α = 0.05. Log Fold Change (LogFC) was calculated using the log-transformed expression data for Dp71 and its variants (log_2_(expected count +1)) as follows: LogFC = mean expression (tumour samples) – mean expression (normal samples).

### Proteomics

Proteins were extracted from BMM on day 7 of differentiation *in vitro*. Macrophages were collected as a pellet, 5-10 times the approximate cell pellet volume of 0.5 M triethyl ammonium bicarbonate (TEAB) with 0.05% SDS was added and cells triturated by passing through a 23-gauge needle 30 times. Samples were then sonicated on ice (30 x 2 sec bursts) and centrifuged at 16000g for 10 min at 4°C. Supernatant was transferred to a fresh Eppendorf tube and protein quantified by nanodrop. 100-150µg of protein was aliquoted for each individual sample and 2µl of 50mM tris-2-carboxymethyl phosphine added for every 20µl of protein. After 1 hr incubation at 60°C, 1µl 200mM methylmethane thiosulphonate) was added for every 20µl of protein. After a 10 min incubation at RT, samples were trypsinised by addition of 6-7.5µl of 500ng/µl trypsin. The ratio between enzyme: substrate was 1: 40. Samples were incubated overnight at 37°C in the dark.

An orthogonal 2D-LC-MS/MS analysis was performed with the Dionex Ultimate 3000 UHPLC system coupled with the ultra-high-resolution nano ESI LTQ-Orbitrap Elite mass spectrometer (Thermo Scientific). An 8-plex iTRAQ labelling and combined CID/HCD fragmentation approach was used. Acquired mass spectra were processed by the Proteome Discoverer 1.4 Software. All PMS were FDR corrected, and validation was based on the q value <0.05. All spectra were searched against the UniProtKB SwissProt murine proteome and quantification ratios were median-normalised. Unified protein and peptide lists were exported from Proteome Discoverer 1.4. Protein ratios were transformed to log_2_-ratios and significant changes were determined by the permutation test followed by LIMMA for bioinformatics analysis with MetaCore software (GeneGo). Significant changes of post-translation modifications ratios (phosphorylated, methylated and acetylated) were determined by two sample t test. Following MetaCore analysis, only pathways with FDR corrected *p<0.05* were considered significantly altered.

#### Cell proliferation assay

Cells were seeded in 10 cm diameter dishes (Sarstedt 83.3902) at 30 % confluency in DMEM F12 with 10% (v/v) FBS, 2mM L- Glutamine, 100U/ml Penicillin and 100U/ml Streptomycin (Sigma) and left to attach for 2 hours before adding BrdU (Invitrogen B23151) to a final concentration of 75 µM and left to proliferate for 6 hours. Cells were then rinsed with warm medium, detached using Accutase (Biowest L0950) and fixed using ice cold ethanol. Samples were left at −20°C overnight and then analysed by flow cytometry using BD FACSCalibur.

#### Cell migration assay

Macrophages in a serum-free medium were seeded on top of the insert (pore size 0.8 µm; Sartstedt 83.3932.800), uncoated or Matrigel coated (Corning 354234), while medium with 10% (v/v) FBS, 2mM L- Glutamine, 100U/ml Penicillin and 100U/ml Streptomycin was placed in the bottom well. Cells were left to penetrate across the filter for 12 hours before fixing in buffered formalin (Sigma HT501128) and staining with Hoechst (1:5000) for cell counting following visualisation under the confocal microscope (LSM 710, Zeiss).

#### Phagocytosis/endocytosis analysis

To analyse phagocytic capability of macrophages, BMM were cultured with fluorescently (CFSE) labelled, IgG-coated sheep red blood cells (SRBC, BIOMED SA Lublin, Poland). Macrophages were seeded in 12-well plates in 0.5ml of media (DMEM F12 with supplements, as described above), at density of 1×10^6^ cells/ml. SRBCs were labelled with 10ìm CFSE (Sigma Aldrich) for 10 minutes at 37°C and washed twice with physiological salt solution. SRBCs were then coated with IgG, by adding 0.05% anti-SRBC serum and incubating for 30 minutes at 4°C. They were then washed 3 times in saline, resuspended in culture media, counted in a haemocytometer and added to BMM at 1:10 macrophage:SRBC ratio. After 1 hour, wells were washed twice with cold PBS to get rid of residual erythrocytes, followed by scraping of macrophages, washing and analysis on either FACSCalibur (BD) or BD LSR Fortessa cytometer.

For dextran uptake analysis, macrophages were seeded in a 6-well plate (Santa Cruz Biotechnology Inc.) at the density of 3 × 10^5^ and were grown in 2% FBS DMEM F12 for 2 h. Cells were then treated with LPS (200ng/ml) or incubated without it for 13 h and then incubated with 1mg/ml of fluorescein isothiocyanate-dextran 40.000kDa (FITC dextran) (Sigma Aldrich Ltd.) for 30 mins, 2 h and 4 h. Two plates were set up for each sample, one incubated at 37°C and another at 4°C. Once incubations were completed, cells were washed twice with ice-cold DPBS, given 2 minutes in HBBS to recover, then removed using ice-cold PBS and cell scrapers. Cells were analysed using either BD FACSCalibur or BD LSR Fortessa cytometer. The plate incubated at 4°C was used as a control to measure the amount of background FITC staining and negative control of dextran uptake.

### Immunolocalization with confocal microscopy

Cells were fixed with 4% (w/v) paraformaldehyde in 1X PBS for 15 min at 4°C and when appropriate, were also permeabilised with <0.1 % Triton X-100 in 1X PBS for 10 min at RT. Cells were washed in 1X PBST (3x) for 5 min after each step. After blocking with 10% normal serum in 1X PBS for 30 min, cells were incubated with primary antibodies (Supplementary Table 3), diluted using 10% (v/v) normal serum from the species that the secondary antibodies were raised in (Vector Laboratories, Burlingame, CA, USA) in 1X PBS at 4°C. Cells were then washed (3x) for 5 min in 1X PBST followed by a 30 min incubation with the respective Alexa Flour secondary antibody (Supplementary Table 3), diluted in 10% (v/v) normal serum in 1X PBS, and Hoechst nucleic acid stain (1:1000). Last wash was performed in 1X PBST (3x) for 10-20 min. Slides were mounted in FluorPreserve™ (Merc Millipore Ltd.) and sealed by applying colourless nail varnish on the edges of the cover slips. Visualisation was achieved using a confocal microscope. Omitting primary antibodies in the reaction sequence served to confirm the specificity of secondary antibodies. Tissue samples were examined with a confocal laser-scanning microscope (LSM 710; Zeiss, Oberkochen, Germany) using either a Plan Apochromat 20x (NA 0.8) (pixel size 0.42 μm) objective, Plan Apochromat 40x DIC oil objective (NA 1.3) (pixel size 0.29 μm), Plan Apochromat 63x DIC oil objective (NA 1.4) (pixel size 0.13 μm) objective or a Plan Apochromat 100x DIC oil objective (NA 1.46) (pixel size 0.08 μm). Images were acquired using sequential acquisition of the different channels to avoid crosstalk between fluorophores. Pinholes were adjusted to 1.0 Airy unit. In all cases where multiple images were captured from the same immunohistochemical reaction, laser power, pinhole, and exposure settings were captured once on tissue from a representative control section and maintained throughout imaging. Images were processed with the software Zen (Zeiss) and exported into bitmap images for processing in Adobe Photoshop (Adobe Systems, San Jose, Ca, USA). Only brightness and contrast were adjusted for the whole frame, and no part of any frame was enhanced or modified in any way.

### Intracellular Ca^2+^ measurements

Peritoneal macrophages were cultured on glass coverslips in 3.5 cm diameter dishes at 70-80 % confluency in the culture medium described above. After 24 h, cells were loaded with Fura-2 AM (Molecular Probes, Oregon) in culture medium for 20 min at 37◦C in a 95% O2, 5% CO2 atmosphere. After two brief washes in the assay buffer (5 mM KCl, 1 mM MgCl2, 0.5 mM Na2HPO4, 25 mM HEPES, 130 mM NaCl, 1 mM pyruvate, 25 mM D-glucose, pH 7.4) the coverslips were mounted in a cuvette and cells were treated with 100 nM thapsigargin (inhibitor of SERCA). The Store Operated Calcium Entry (SOCE) was measured in fluorescence spectrophotometer (F-7000 Hitachi) after adding 2mM calcium to the assay buffer at room temperature. Fluorescence was recorded at 510nm with excitation at 340/380 nm. Delta ratio of both signals was then calculated. Each experiment was repeated three times.

### Statistical analysis

Results are reported as mean ± SD, where *n* refers to the number of independent biological replicates. Unpaired Student t-test was used for comparisons between the two data groups and one-way ANOVA with Tukey post-hoc analysis (Minitab 17) when more than two groups were compared. For proteomics analysis, LIMMA following permutation test (R environment) was applied for the protein list and two sample T test was performed for phospho, acetylated and methylated protein lists, *p**-***value of <0.05 was considered statistically significant, and the values are reported as follows: \**p **<*** 0.05, \*\**p **<** 0.005*, \*\*\**p < 0.001*.

## Acknowledgements

We thank Michael Nemeth for his helpful suggestions on the HSC analysis, K Zablocki for the critical comments on the manuscript.

## Author’s contributions

DCG designed research; NC, JS, AM MRFG, JH, CY, AO, DGB, JR, CHW, SA, MC, EK, NA, SDG and DCG performed research; JH, MK, EK, SDG contributed reagents/analytic tools and SDG supervised the proteomic analyses; NC, JS, AM, MRFG, JH, CY, AO, DGB, JR, CHW, SA, MC, EK, SDG and DCG analysed data; NC and DCG wrote the draft paper and all authors contributed to its final version.

