## Supplementary figures for "Dystrophin Dp71 is essential for the development and function of macrophages"

**Loss of DMD gene expression triggers cell-autonomous abnormalities in the functional development of macrophages.**

### Supplementary Figures

A

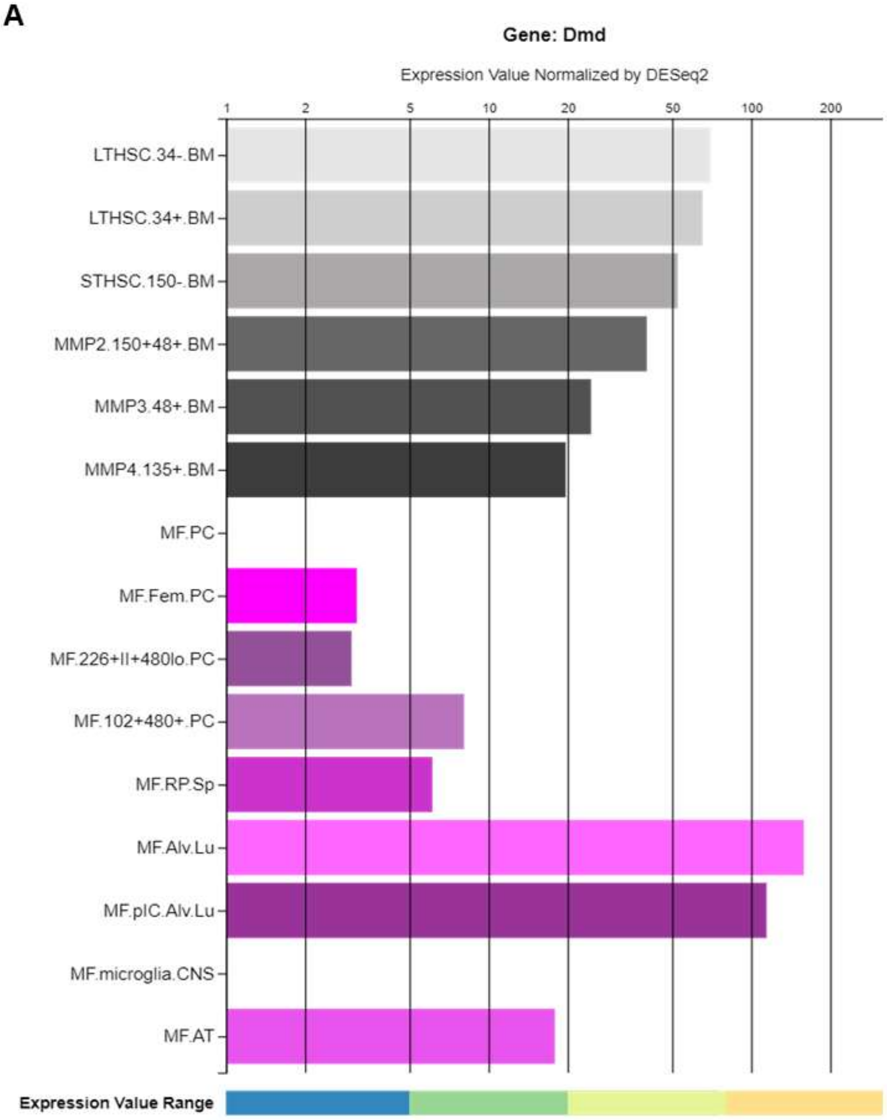


B

| Cell | HSC | MPP1 | MPP2 | MPP3 | MPP4 |
|---|---|---|---|---|---|
| ENSMUST00000114000 (dp427) rank | 67009 | 61291 | 62829 | 67634 | 70037 |
| ENSMUST00000239019 (dp71) rank | 23765 | 20124 | 24930 | 34388 | 33596 |
| n = number of analyzed transcripts | 142526 | 142526 | 142526 | 142526 | 142526 |
| Dp427 / Hprt [%] | 0.01 | 0.02 | 0.02 | 0.01 | 0.01 |
| Dp427 / Gapdh [%] | 0.04 | 0.05 | 0.08 | 0.04 | 0.03 |
| Dp71 / Hprt [%] | 0.82 | 0.83 | 0.46 | 0.26 | 0.28 |
| Dp71 / Gapdh [%] | 2.33 | 2.62 | 2.19 | 0.88 | 0.96 |

**Supplementary Fig. 1.**

DMD gene expression in differentiating HSCs and developed macrophages of different origins. (**A**) *Dmd* gene expression values in HSC and various stages of lineage differentiation compared to macrophages. Data obtained from the Immunological Genome Project database (https://www.immgen.org/) using the Gene Skyline functionality (http://rstats.immgen.org/Skyline/skyline.html). Cell descriptions:

LTHSC.34-.BM Bone Marrow 34- LTHSC/Bone Marrow 34- Long Term hematopoietic stem cells from 6-week-old C57BL/6J mice sorted on Lin-Sca1+ckit+CD135-CD150+CD48-CD34-.

LTHSC.34+.BM Bone Marrow 34+ LTHSC/Bone Marrow 34+ Long Term hematopoietic stem cells from 6-week-old C57BL/6J mice sorted on Lin-Sca1+ckit+CD135-CD150+CD48-CD34+.

STHSC.150-.BM Bone Marrow CD150- Short Term HSC/Bone Marrow CD150- Short Term Hematopoietic Stem Cell from 6-week-old C57BL/6J mice sorted on Lin-Sca1+ckit+CD135-CD150-CD48-

MPP2.150+48+.BM Bone Marrow CD150+ CD48+ MMP2/Bone Marrow CD150+ CD48+.

MPP3.48+.BM Bone Marrow CD150+ CD48+ MMP3/Bone Marrow CD150+ CD48+, from 6-week-old C57BL/6J mice sorted on Lin-Sca1+ckit+CD135-CD150-CD48+.

MPP4.135+ BM Bone Marrow CD150+ CD48+ MMP4/Bone Marrow CD150+ CD48+, from 6-week-old C57BL/6J mice sorted on Lin-Sca1+ckit+CD135+.

MF.PC. Peritoneal Macrophages, 6-week-old C57BL/6J mice sorted on F4/80+ICAM2+CD5-CD19-CD43-.

MF.Fem.PC. Female Peritoneal Macrophages from 6-week-old C57BL/6J mice sorted on F4/80+ICAM2+CD5-CD19-CD43-.

MF.226+II+480lo.PC. Peritoneal Small Macrophages from 6-week-old C57BL/6J mice sorted on CD115+CD11b+F4/80loCD102loMHCII+CD226+.

MF.102+480+.PC. Peritoneal Large Macrophages, 6-week-old C57BL/6J mice sorted on CD115+CD11b+F4/80+CD102+MHCIIloCD226-

MF.RP.Sp. Red Pulp Macrophages, 6-week-old C57BL/6J mice sorted on Mertk+ CD64+ CD11blo F4/80+

MF.Alv.Lu. Alveolar Macrophages, 6-week-old C57BL/6J mice sorted on CD45+ CD11c+ SiglecF+.

MF.pIC.Alv.Lu. Alveolar Macrophages, Poly IC, 6-week-old C57BL/6J mice sorted on CD45+ CD11c+ SiglecF+.

MF.microglia.CNS. Brain Microglia Macrophages, 6-week-old C57BL/6J mice sorted on CD19-CD3-Ly6G-CD45loCD11b+ CD64+ F4/80loCD206-.

MF.AT. Inguinal & Perigonadal Adipose Tissue Macrophages, 6-week-old C57BL/6J mice sorted on CD19-Ly6G- CD45+CD11b+F4/80+MerTK+CD64+.

Note, while this analysis confirmed the decreased *Dmd* expression in BMM and PMφ, in alveolar macrophages it was still significant, possibly reflecting their different developmental origin from the yoke sac[100,101].

(**B**). DMD gene expression at different stages of HSC development. Table represents rankings of specific *Dmd* gene transcripts amongst the 142.526 transcripts expressed in these cells, as identified in [49] as well as the relative expression of these *Dmd* transcripts (%) against *Hprt* and *Gapdh* housekeeping genes.

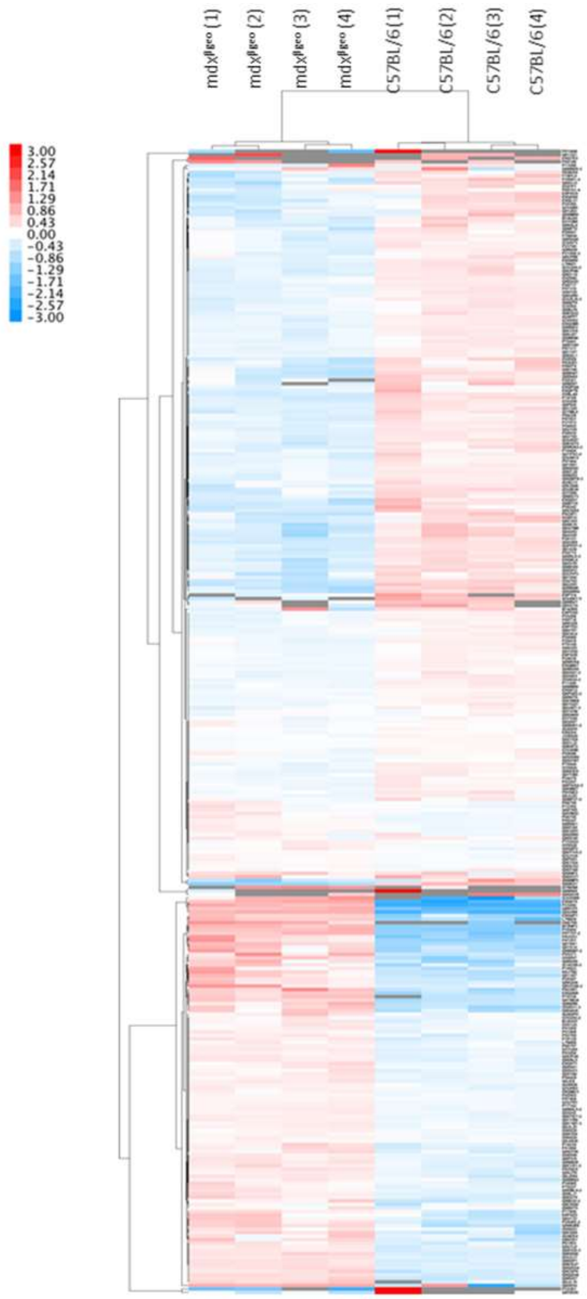

**Supplementary Fig. 2.** Heatmap based on the hierarchical clustering of differentially expressed proteins in mdx$^{βgeo}$ vs. C57BL/6 BMM. This heat map illustrates the global trends. It is not intended for the analysis of individual proteins. These are presented in the Supplementary Table 1.

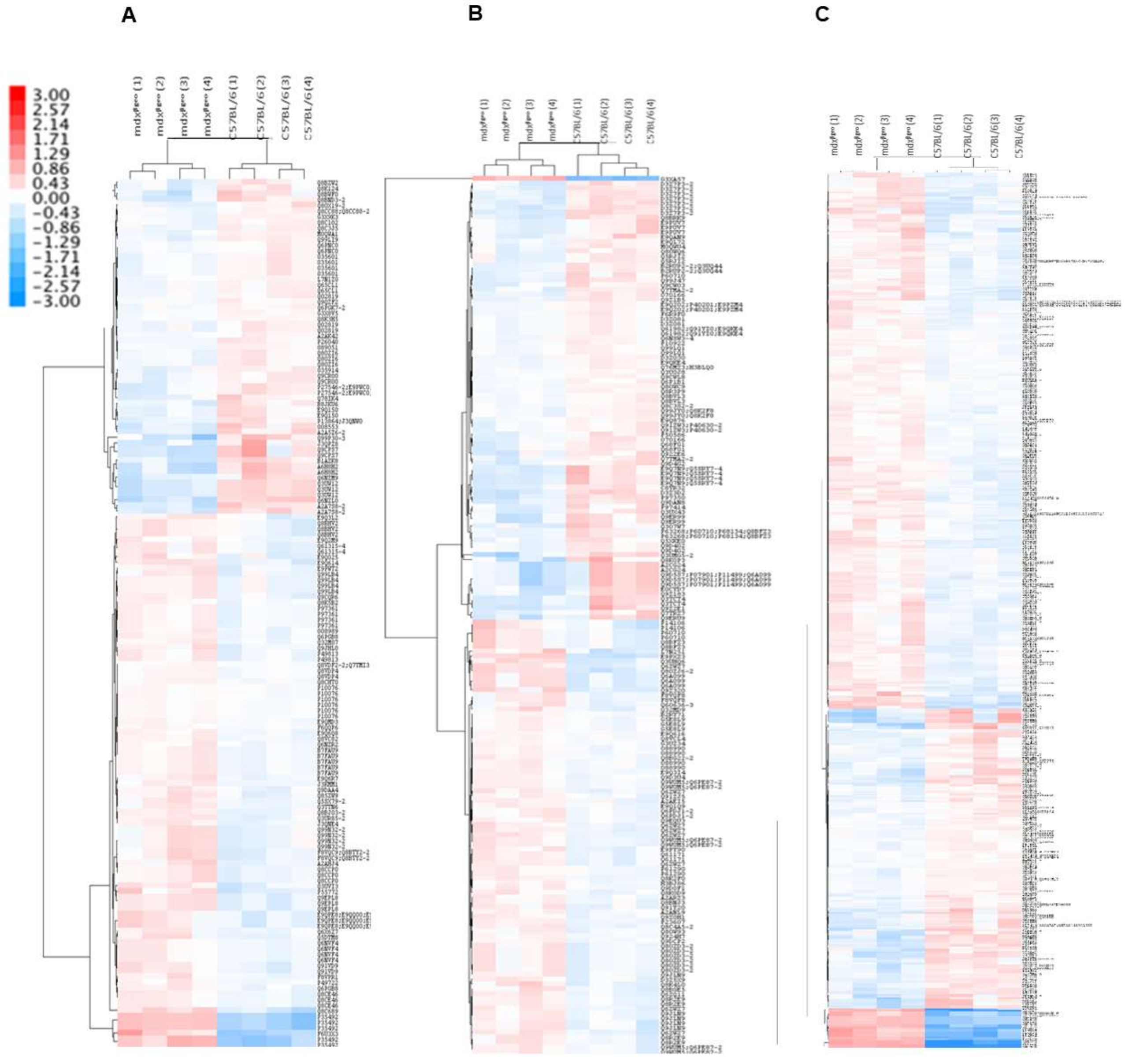


**Supplementary Fig. 3.** Heatmap based on the hierarchical clustering of differentially expressed: (**A**) phospho-proteins, (**B**) acetylated proteins and (**C**) methylated proteins in mdx$^{βgeo}$ vs. C57BL/6 BMM. These heat maps illustrate global trends and are not meant for the analysis of individual proteins.

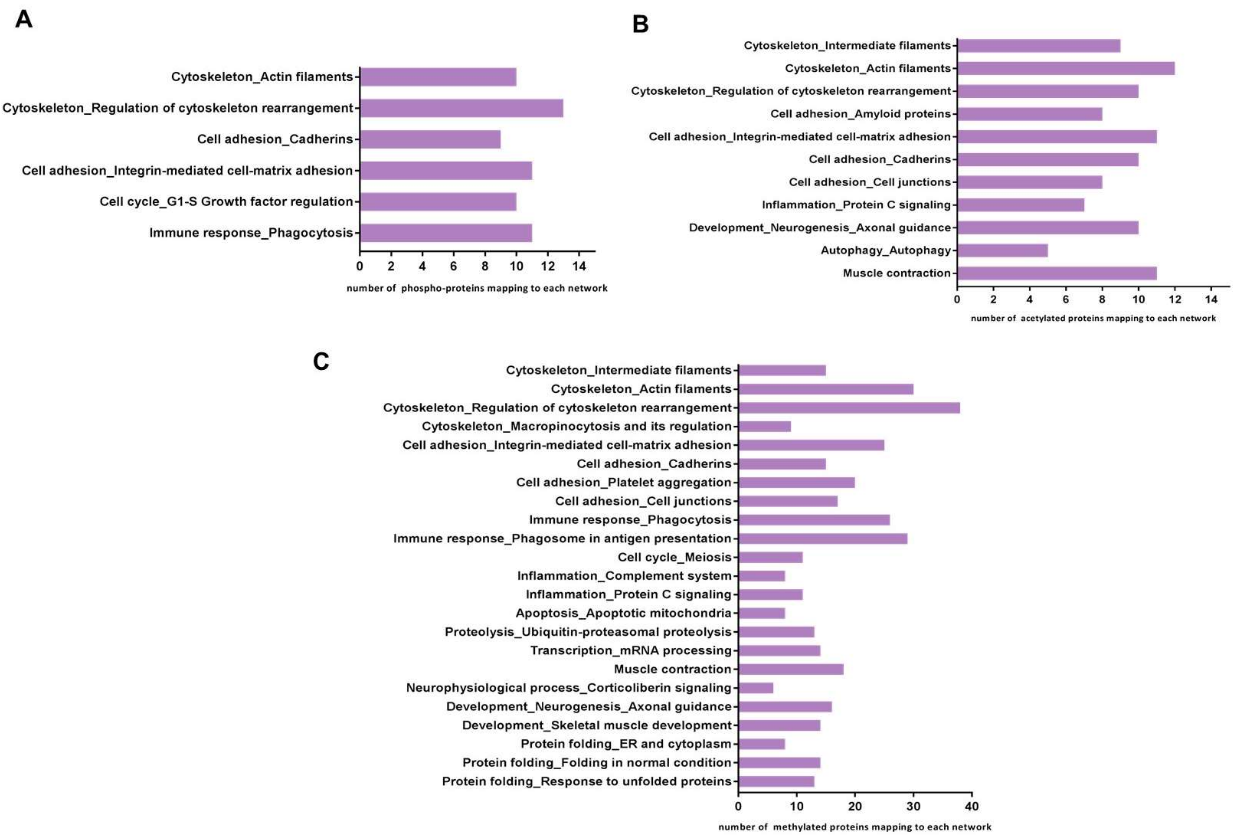


**Supplementary Fig. 4.** Differentially expressed process networks/pathways between wild-type and $mdx^{\beta geo}$ BMMs. MetaCore was used to map identified proteins onto the defined networks, based on the known protein–protein interactions and other features established in the literature. Enrichment analysis of pathways based on **(A)** the phospho-protein list, **(B)** the acetylated-protein list and **(C)** the methylated-protein list.

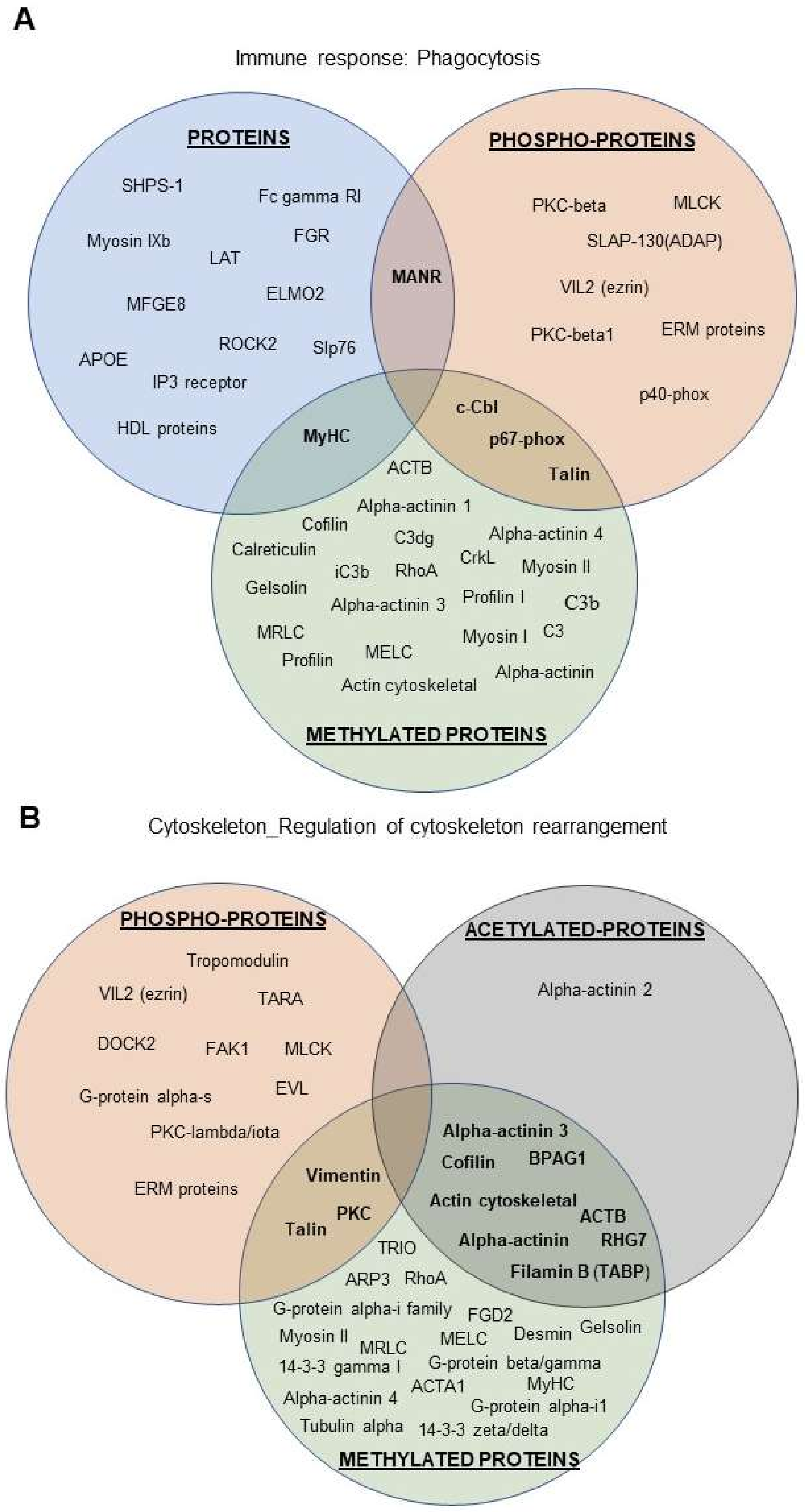


**Supplementary Fig. 5.** Venn diagrams demonstrating common alterations identified between the total protein, phosphoprotein, acetylated and methylated protein datasets for the immune response_phagocytosis and cytoskeleton_regulation of cytoskeleton rearrangement networks.

| Protein name | log2FC mdx$^{\beta geo}$ / C57BL/6 |
|---|---|
| **Macrophages markers** | |
| CD45 | n.s |
| MCSF receptor factor 1 (CD115) | n.s |
| FcγRI (CD64) | -0.91 |
| MerTK | n.s |
| F4/80 | n.s |
| Short isoform of CD68 | n.s |
| Fert2 | n.s |
| **Proteins related to M1 polarisation** | |
| GM CSF receptor | 0.38 |
| IL-12b | n.s |
| IL-6 | n.s |
| IL-18 | n.s |
| VCAM1 | n.s |
| CD80 | n.s |
| CD86 | n.s |
| **Proteins related to M2 polarisation** | |
| Arg | n.s |
| CD206 | -0.69 |
| TGFB1 | n.s |
| Maf | -0.47 |
| PPARγ | n.s |
| CHI3L3 (YM-1) | n.s |
| **Phagocytosis related proteins** | |
| Gas6 | -0.74 |
| MFGE8 | -0.52 |
| Axl protein | n.s |
| FcγRI | -0.91 |
| CD206 | -0.69 |
| MerTK | n.s |

**Supplementary Fig. 6**. Comparison of macrophage markers in wild-type and mdx$^{\beta geo}$ BMM. Comparison of the expression of M1/M2 polarisation markers, opsonins and receptors related to the phagocytic activity in MS data from the wild-type and mdx$^{\beta geo}$ macrophages. Blue indicates downregulation and red upregulation of the protein expression in mdx$^{\beta geo}$ compared to wild-type BMM, respectively. n.s - no significant differences.

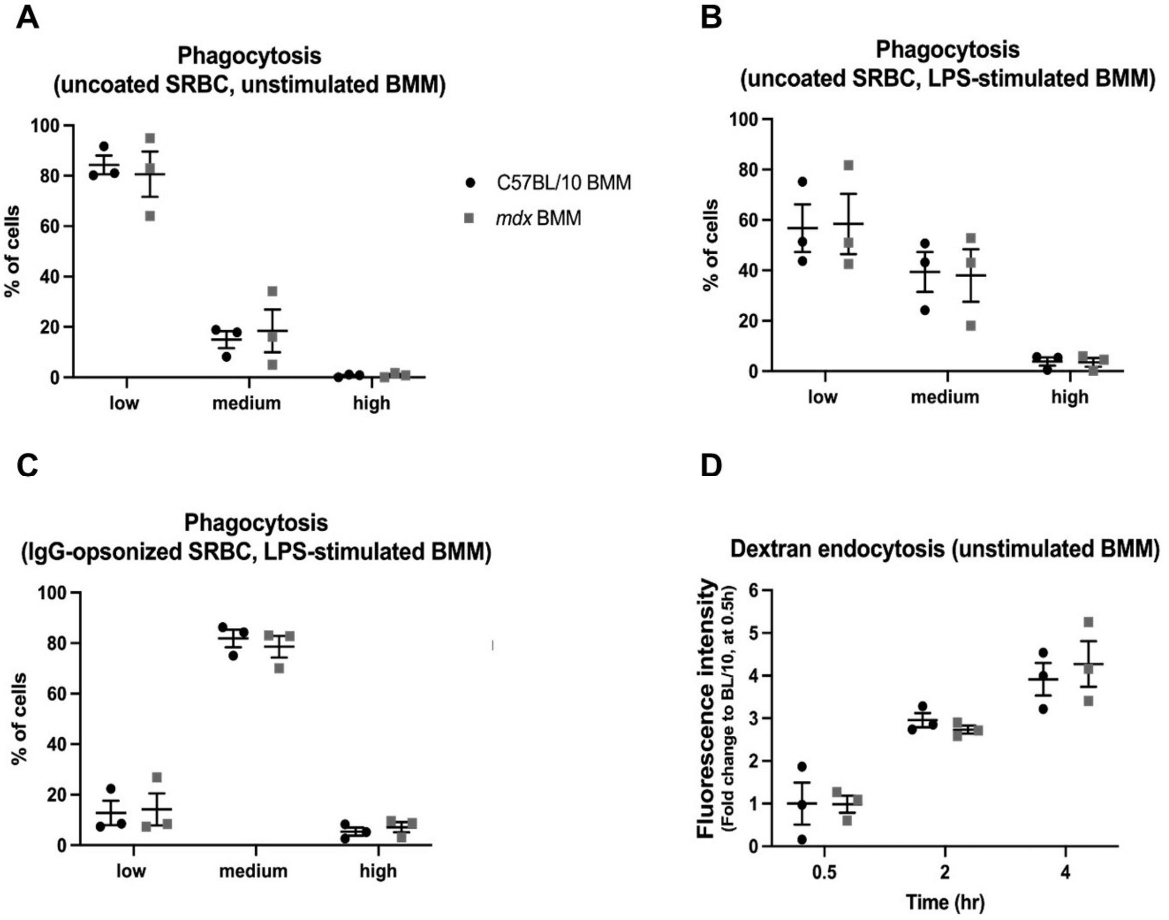


**Supplementary Fig. 7.** Functional assays of endocytic activity of mdx BMM. **(A)** Phagocytosis of un-opsonized sheep red blood cells (SRBCs) by unstimulated BMM. (**B**) Phagocytosis of un-opsonized SRBCs and (**C**) Fc mediated phagocytosis of IgG-opsonized SRBCs by LPS-stimulated BMM. In (A)-(C), percentage (%) of cells that were gated as non-/low, medium or highly phagocytic are shown on the graphs as single data points, mean $\pm$ SEM (unpaired t-test with Welch's correlation). **(D)** Endocytosis of dextran-FITC (40.000 MW) particles by unstimulated BMM. Fluorescence intensity (corresponding to dextran uptake, values normalized to WT C57BL/6 BMM at 0.5h in each experiment) in cells, following 0.5, 2, and 4-hour incubation with dextran, is shown on the graphs as single data points, mean $\pm$ SEM (unpaired t-test with Welch's correlation).

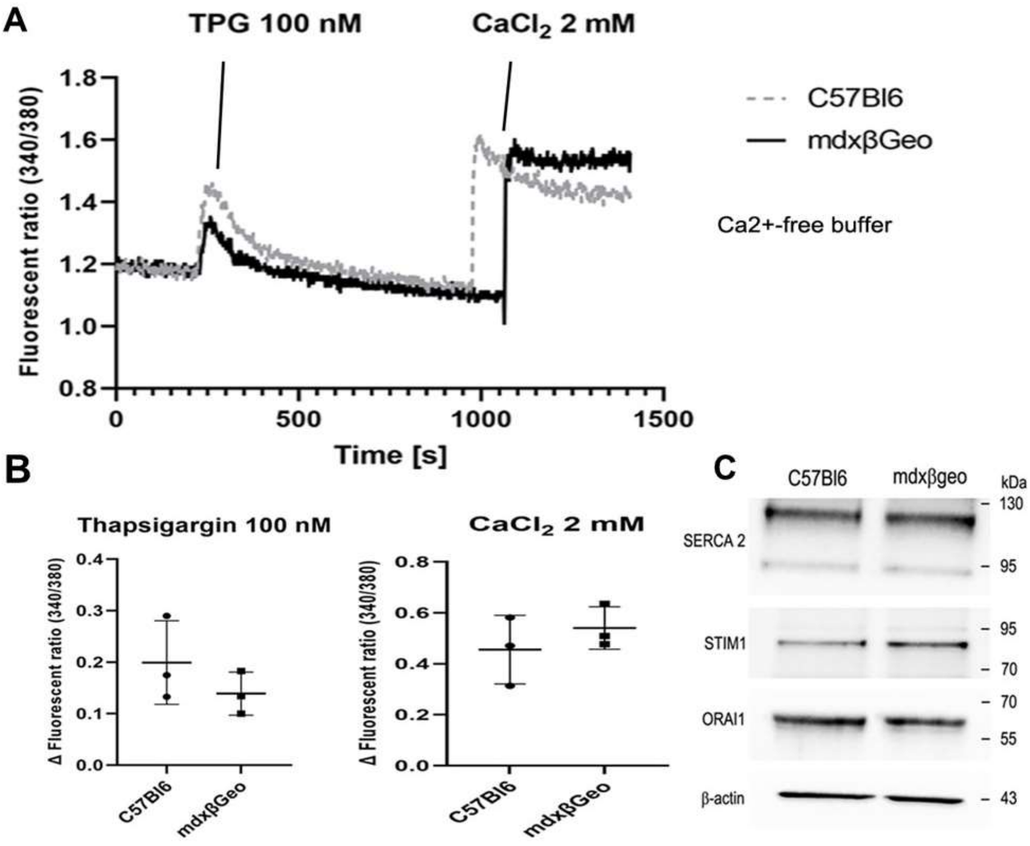


**Supplementary Fig. 8.** Analysis of the store-operated calcium entry (SOCE) and expression of relevant proteins in primary peritoneal macrophages.

(**A**) Representative traces from measurements of changes in cytosolic $Ca^{2+}$ concentrations using Fura 2AM dye expressed as a ratio of fluorescence at the excitation wavelength 340 and 380 nm. Cells in $Ca^{2+}$ free buffer were treated with 100 nM thapsigargin to empty the ER calcium stores (the first peak). Subsequent addition of 2 mM $CaCl_2$ initiates $Ca^{2+}$ entry and leads to the fast elevation of the cytosolic $Ca^{2+}$ concentration (the second peak) due to activated SOCE. (**B**) Summary data from three independent experiments performed as described above. No significant difference in SOCE between C57BL/6 and $mdx^{\beta geo}$ PMϕ. (**C**) Representative Western blot images of SERCA2, STIM1 and ORAI1 in cell lysates from C57BL/6 and $mdx^{\beta geo}$ PMϕ showing no significant differences in the expression levels of these proteins. β-actin used as a protein loading control.

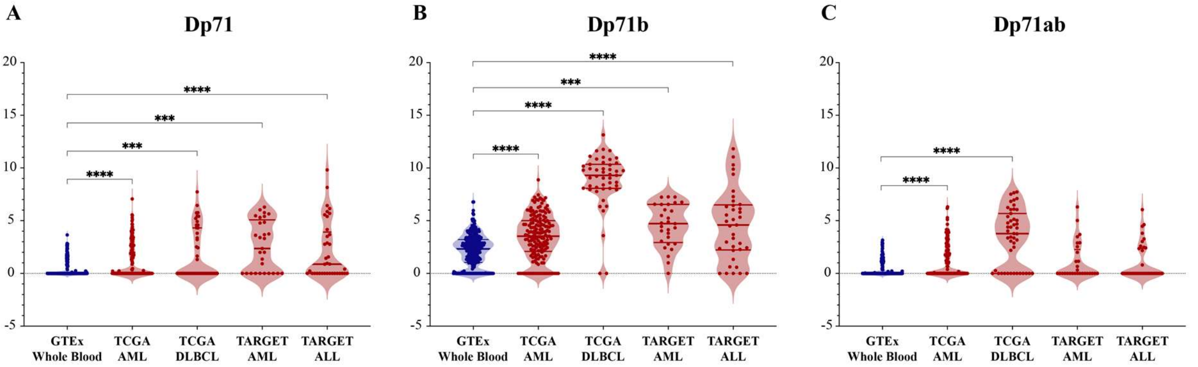


**Supplementary Fig 9.** Expression of [**A**] Dp71, and its specific splice variants [**B**] Dp71b (missing exon 78) and [**C**] Dp71ab (missing exons 71 and 78) in primary blood malignancies compared to whole blood. Blue violin plots represent healthy whole blood from the GTEx database, and red violin plots represent blood malignancies: acute myeloid leukaemia (AML) and diffuse large B-cell lymphoma (DLBCL) from the TCGA database, and AML and acute lymphoblastic leukaemia (ALL) from the TARGET database. Vertical lines represent the median and quartiles. P-values were calculated using the Xena Browser using a two-tailed Welch's t test and adjusted for multiple testing using the Bonferroni correction (***p < 0.001, ****p < 0.0001).

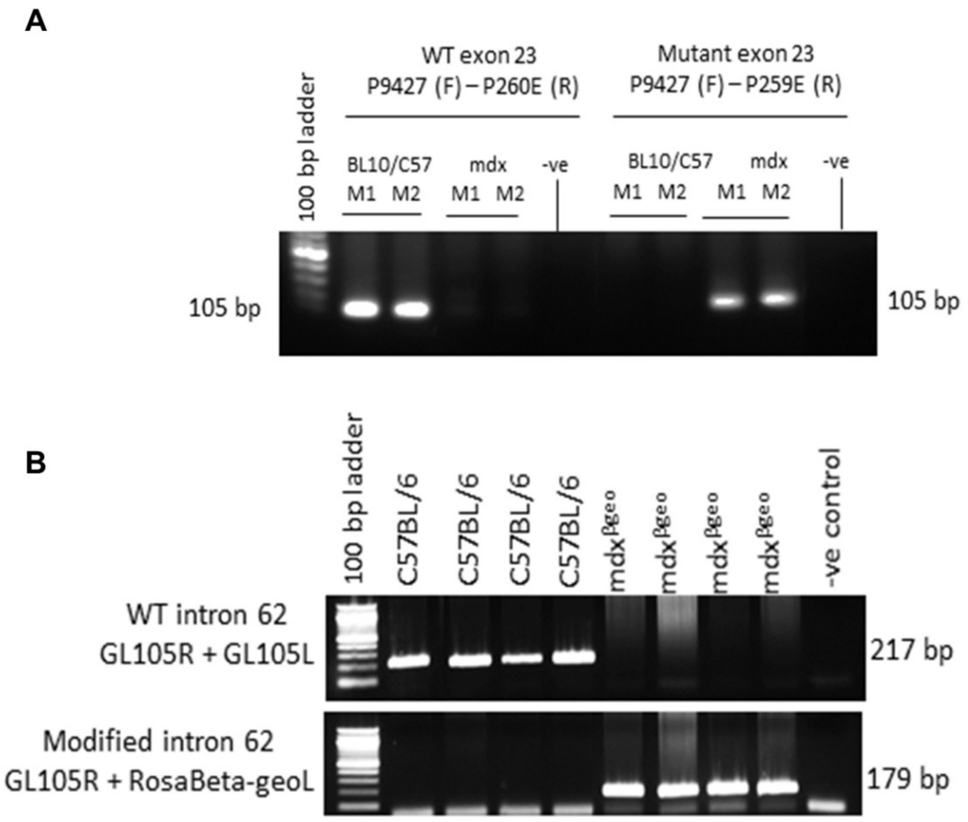

**Supplementary Fig. 10.** Genotyping of experimental animals. Example images of gel electrophoresis of PCR amplicons confirming the genotypes of experimental animals. **(A)** C57BL/10 samples showing the expected 105bp band with the primer set complementary to the wild type exon 23 and mdx samples having a 105bp band amplified with primer set complementary to the mutant exon 23 of the *Dmd* gene. **(B)** C57BL/6 samples showing amplification of the expected 217bp band with a primer set complementary to wild type intron 62, and in mdx$^{\beta geo}$ samples amplification of a 179bp band with a primer set complementary to the modified intron 62 of the *Dmd* gene confirmed the expected genotype. -ve control = a negative control where gDNA sample was replaced with water. M1, M2 denote the individual mouse numbers.
